# Single nucleus and spatial transcriptomic profiling of human healthy hamstring tendon

**DOI:** 10.1101/2022.12.19.521110

**Authors:** Jolet Y. Mimpen, Lorenzo Ramos-Mucci, Claudia Paul, Alina Kurjan, Phillipa Hulley, Chinemerem Ikwuanusi, Steve Gwilym, Mathew J. Baldwin, Adam P. Cribbs, Sarah J.B. Snelling

## Abstract

The molecular and cellular basis of health in human tendons remains poorly understood. Amongst human tendons, the hamstrings are the least likely to be injured or degenerate, providing a prototypic healthy tendon reference. The aim of this study was to define the transcriptome and location of all cell types in healthy hamstring tendon. We profiled the transcriptomes of 10,533 nuclei from 4 healthy donors using single-nucleus RNA sequencing (snRNA-seq) and identified 12 distinct cell types. We confirmed the presence of two fibroblast cell types, endothelial cells, mural cells, and immune cells, and revealed the presence of cell types previously unreported for tendon sites, including different skeletal muscle cell types, satellite cells, adipocytes, and nerve cells, which are undefined nervous system cells. Location of these cell types within tendon was defined using spatial transcriptomics and imaging, and transcriptional networks and cell-cell interactions were identified. We demonstrate that fibroblasts have a high number of potential cell-cell interactions, are present throughout the whole tendon tissue, and play an important role in the production and organisation of extracellular matrix, thus confirming their role as key regulators of hamstring tendon tissue homeostasis. Overall, our findings highlight the highly complex cellular networks underpinning tendon function and underpins the importance of fibroblasts as key regulators of hamstring tendon tissue homeostasis.

## 1 Introduction

Tendons are fibrous connective tissues that seamlessly connect muscles to bone. The fibrous composition of the tendons allows them to withstand large mechanical loads, and store and deliver substantial forces that facilitate joint movement and stability[1]. Despite their important role in musculoskeletal function, there is still a lack of understanding of the underlying biology of tendon; even the cellular composition and prominent markers remain poorly understood compared to neighbouring musculoskeletal tissues. Consequently, treatment of tendon disease pathology such as tendinopathy is limited to exercise programmes, pain management, and surgical intervention, but largely fails to address disease pathology or allow focussed drug development[2].

The tendon extracellular matrix (ECM) is mainly made up of type I collagen, type III collagen and proteoglycans, including decorin, cartilage oligomeric matrix protein, fibromodulin, and biglycan, among others[3]. ECM components are produced and maintained by fibroblast-like cells, also called tenocytes. These elongated fibroblasts are located inside collagen fibres and in the surrounding endotenon, the loose connective tissue which surrounds collagen bundles in tendons. The endotenon contains a more heterogeneous mix of vascular, nerve, and mesenchymal cells[4]. Furthermore, the presence of both innate immune cells, such as macrophages, and adaptive immune cell types, such as T cells, have been reported[5]. These studies demonstrate a diverse cellular environment, yet these cell types and their interactions have not been comprehensively characterised. Insight into the cellular composition of healthy tendon is vital to unravel pathways and relationships dysregulated in disease, enabling the identification of new pharmaceutical targets and better design of biomaterials. Currently, progress in the field of tendon biology is hindered by accessibility of healthy comparative tissue and the technical challenges posed by ECM-rich tissues[6, 7].

Single-cell RNA sequencing (scRNA-seq) technologies are increasingly used to characterise major and rare cell populations within tissues and infer important cellular markers and potential cellular interactions. Initial studies by Kendal *et al*. (2020) and Akbar *et al*. (2021) presented CITE-seq and scRNA-seq results of healthy and diseased tendons. Despite the limited number of cells captured in these studies, ^~^440 cells and 3,040 cells, respectively [8, 9], identification of fibroblasts/stromal cells, immune cells, and endothelial cells was possible. However, the low cell numbers captured limited a detailed cellular profiling of healthy tendon. Therefore, a more comprehensive and spatially resolved dataset is needed to infer the cellular states, functions, and interactions that underlie hamstring tendon health. The aim of this study was to create a comprehensive single-cell healthy hamstring reference dataset using single-nucleus RNA-sequencing (snRNA-seq), and resolve cellular locations using spatial transcriptomics and immunofluorescence imaging.

## 2 Methods

### 2.1 Ethics

Ethical approval was granted for the Oxford Musculoskeletal Biobank (19/SC/0134) by the local research ethics committee (Oxford Research Ethics Committee B) for all work on human hamstring tendon, and informed written consent was obtained from all patients according to the Declaration of Helsinki.

### 2.2 Tissue acquisition and processing

Healthy hamstring tissue (semitendinosus; proximal one-third which includes the midbody to the myotendinous junction) was collected from patients undergoing ACL reconstruction for an ACL rupture, in which healthy hamstring is used as a graft tissue. The age, gender, affected side, and body mass index were collected (Table 1). Tendon was handled as previously described [11]. Tissue was collected in cold DMEM-F12 media supplemented with 10% Foetal Bovine Serum and 1% Penicillin/Streptomycin and transferred to the research facility for processing. Within two hours of tissue collection, tendon was washed in PBS, fat and muscle were cut off, and the tendon was cut into 1 cm pieces and photographed to retain topographical reference. All the pieces were snap-frozen in cryotubes using liquid nitrogen. Tissue was stored at −80°C until use.

**Table 1.** Patient characteristics, including sex, age, body-mass index (BMI), affected side, and ethnicity.

|  | Sex | Age (yrs) | BMI | Side | Ethnicity | Single nucleus RNA-Seq | Spatial transcriptomics | Imaging |
| --- | --- | --- | --- | --- | --- | --- | --- | --- |
| Hamstring 0 | Male | 37 | 24.38 | Right | Unknown | Yes | No | No |
| Hamstring 1 | Male | 24 | 27.78 | Right | White – British | Yes | No | No |
| Hamstring 2 | Female | 26 | Unknown | Right | White – British | Yes | No | No |
| Hamstring 5 | Female | 18 | 29.73 | Left | White – British | Yes | Yes | Yes |

### 2.3 Single-nucleus RNA sequencing

#### 2.3.1 Nuclei isolation

Nuclei were isolated using our previously published protocol[12]. In short, forceps, scalpels, and petri dishes were pre-cooled on dry ice. Tissue was cut into thin flakes and stored in a pre-cooled 50 mL Falcon tube. The tissue was then stored at −80°C until use. On the day of cell lysis, the tubes containing the pre-cut tissue were thawed and 4mL of cold 1X CST buffer (salts and Tris buffer, including NaCl, Tris-HCl pH 7.5, CaCl2, and MgCl2, with CHAPS hydrate (Sigma), BSA (Sigma), RNase inhibitors (RNaseIn Plus (Promega) and SUPERase In (Invitrogen)), and protease inhibitor (cOmplete tablet, Roche); full recipe can be found in the published protocol) was added. After 10 minutes of incubation on a rotor at 4°C, the tissue/buffer mixture was poured through a 40 μm strainer and the tube used for tissue lysis was washed twice with 2mL PBS with 1% BSA. The nuclei solution was then transferred to a 15mL Falcon tube, and the previous 50mL tube was washed once with 4 mL PBS with 1% BSA. The solution was centrifuged at 500g at 4°C for 5 minutes. After pouring off the supernatant, the tubes were briefly spun down to get all the liquid to the bottom. The nuclei were then resuspended in the remaining supernatant, and remaining volume was determined. Concentration of nuclei was determined by staining the nuclei with DAPI and counting using a Neubauer Improved haemocytometer (NanoEnTek).

#### 2.3.2 Library preparation and sequencing

Nuclei suspensions were diluted (PBS with 1% BSA) to 200-1000 nuclei/μl and loaded on the Chromium Next GEM Chip G (10x Genomics) with a targeted nuclei recovery of 5,000-10,000 nuclei per sample. Samples were then loaded to the Chromium Controller (10x Genomics) and libraries were prepared using the Chromium Next GEM Single Cell 3’ Reagent Kits v3.1 (10x Genomics) following the manufacturer’s instructions and indexed with the single-index kit T Set A (10x Genomics). Quality control of cDNA and final libraries was analysed using High Sensitivity ScreenTape assays on a 4150 TapeStation System (Agilent). Final libraries were pooled and sequenced on a NovaSeq 6000 (Illumina) by Genewiz (UK) at a minimum depth of ^~^20,000 read pairs per expected nuclei.

#### 2.3.3 Single nucleus RNA-seq data analysis

Raw NGS data was processed using an in-house pipeline scflow quantnuclei (https://github.com/cribbslab/scflow). Quality control of Fastq files was performed using fastqc. Fastq files were mapped to the human genome hg38 using kallisto bustools kb count (release 99) with kmer size = 31 (https://www.kallistobus.tools/kb_usage/kb_count/). Spliced and unspliced matrices were merged to create the count matrix for downstream analysis.

The snRNA-seq analysis was performed using Seurat v4.0 [15]. Filtering thresholds for number of cells (nCount, number of features(nFeature) and mitochondrial ratio (mitoRatio) were set manually for each sample to remove poor-quality cells (Table 2). Doublets were detected and removed with scDblFinder using the default settings[13]. Ambient RNA (score > 0.2) was removed using decontX from the package celda [14] (Supplementary Figure 1). After quality control, samples were merged, normalised (based on 2000 variable features) and scaled. Principal Component Analysis (PCA) was performed and the first two PCs were visualised to assess variation in sample, batch, sex, and side (Supplementary Figure 2). All four samples were integrated by calling Harmony from within the Seurat workflow, using sample as the covariate [16]. Dimensionality reduction was performed on the integrated data using UMAP and t-SNE, neighbours were identified, and clustering was performed (50 dimensions, resolution 0.15; Supplementary Figure 3). Clustering was visualised on UMAP to assess variation in sample, batch, sex, or side (Supplementary Figure 4). Differential expression analysis was performed using FindMarkers (min.pct = 0.25) and the top 5 positive DE genes were visualised using pheatmap (https://www.rdocumentation.org/packages/pheatmap/versions/1.0.12/topics/pheatmap). Dotplots, volcano plots and violin plots were generated using ggplot2. Pathway analysis was performed using gprofiler2[17] to identify over-represented pathways from Gene ontology biological processes (GO:BP) and Reactome databases using the top 100 differentially expressed genes for each clusters (log2 fold change > 0.5). R scripts are available at https://github.com/Botnar-MSK-Atlas/hamstring_atlas.

**Table 2.** Quality control of snRNA-Seq samples, including the number of nuclei before filtering, the minimum and maximum number of Features (nFeature) and number of Counts (nCount), the maximum mitochondrial ratio (mitoRatio), and the number of nuclei after filtering and doublet and ambient RNA removal.

|  | Nuclei before filtering (n) | nFeature minimum | nFeature maximum | nCount minimum | nCount maximum | mitoRatio maximum | Final nuclei (n) |
| --- | --- | --- | --- | --- | --- | --- | --- |
| Hamstring 0 | 5,378 | 800 | 3,000 | 1,000 | 13,000 | 0.05 | 4,301 |
| Hamstring 1 | 5,366 | 500 | 3,000 | 1,000 | 13,000 | 0.05 | 4,653 |
| Hamstring 2 | 249 | 500 | 3,000 | 1,000 | 13,000 | 0.05 | 120 |
| Hamstring 5 | 1,985 | 500 | 3,000 | 1,000 | 10,000 | 0.025 | 1,459 |
| Total | 12,978 |  |  |  |  |  | 10,533 |

#### 2.3.4 SCENIC analysis

Analysis was carried out on the command line and in a Jupyter Notebook in an anaconda environment with Python v3.7.12 and pyscenic v0.11.2, among other packages. Briefly, harmony-integrated Seurat data was converted into the h5ad format using Sceasy[18]. The raw, unnormalized counts matrix as well as the accompanying anndata observations and variables were made into a new object and saved as a .loom file, which was used as input for the pySCENIC workflow steps[19, 20].

Gene regulatory network (GRN) inference was carried out using the GRNBoost2 algorithm by running the “pyscenic grn” command using default settings. A predefined list of human transcription factors used for this network inference step was retrieved from the Aertslab github repository (https://github.com/aertslab/SCENICprotocol/tree/master/example, ‘allTFs_hg38.txt’, last updated 3 years ago). The resulting list of TF-gene interactions was then used to infer TF-gene co-expression modules, identify enriched motifs and predict regulons by running the “pyscenic ctx” command with default settings. Input genome rankings .feather and motif annotation .tbl files were retrieved from the Aertslab cistarget resources webpage (v9 files; https://resources.aertslab.org/cistarget/). In total, 317 regulons were identified. Next, the activity of predicted regulons in individual cells was quantified by running the “pyscenic aucell” command using default settings. For each regulon, cellular regulon activity was then binarized as “on” or “off” using a Gaussian mixture model. Additionally, we assessed regulon-cell type cluster specificity using the regulon specificity score (RSS).

#### 2.3.5 CellPhoneDB analysis

Single-nuclei analysis of the interactions between different cells were analysed using cellphoneDB v3.1.0 (https://github.com/ventolab/CellphoneDB)[21]. A table of the metadata and the expression matrices were exported from the integrated and annotated hamstring Seurat object. For statistical analysis, receptors and ligands expressed in more than 10% of nuclei in a specific cluster were considered. Further analysis of CellPhoneDB predictions, including biological functions and pathways associated with ligand and receptor genes, was completed with InterCellar (v2.0.0)[22].

### 2.4 Spatial transcriptomics (Visium)

Human hamstring tendon from one of the patients analysed for snRNA-seq was also taken for spatial transcriptomics (see Table 1). Snap frozen tissue was embedded in cold OCT mounting medium (VWR) on dry ice. Frozen sections were cut at 10μm thickness through the coronal plane of the tendon and mounted on a 6.5 × 6.5mm capture area of a Visium spatially barcoded slide (Visium Spatial Gene Expression, 10x Genomics). Following the manufacturer’s recommendations, sections were fixed, stained with H&E, imaged, permeabilised and reverse transcribed. cDNA was then released, collected, and prepared for sequencing. Libraries were sequenced on an Illumina Nextseq 500 at a depth of 48,217 mean reads per spot. Data and image alignment was processed using SpaceRanger (v.1.2.2, 10X Genomics). Feature-barcode matrices and H&E images were then used for downstream analysis.

Downstream analysis was conducted using Seurat [15], STUtility. Briefly with STUtility, samples filtered barcodes were loaded and filtered (>20 features/spot and <25% mitochondrial reads) remaining with 2694 spots after filtering. Data was normalised using SCTransform[23] and features of interest were visualised using FeatureOverlay.

Visium data allowed for visualisation of cell types previously identified in single-nuclei transcriptomics using Cell2location[24, 25]. Cell2location estimates the abundance of cell types in each Visium spot by integrating the two datasets. For this analysis, the annotated single-nuclei dataset (Seurat object) was converted to h5ad using Sceasy and used as the reference dataset. First, the reference expression signatures for all cell types (n=12) are estimated using a negative binomial regression which accounted for donor effects. Second, the reference signatures were used to estimate absolute abundance in the Visium spots with the following hyperparameters (cell abundance per spot = 3 and detection alpha = 10). Cell abundance was estimated using histology and detection alpha was kept low to allow for more lenient regularisation to account for large technical variability in RNA content. The estimated cell abundance was plotted for each cell type or multiple cell types in the Visium slide.

### 2.5 Immunofluorescence staining and imaging using Cell DIVE

Snap-frozen tendon samples were embedded in OCT, before cutting 7 mm tissue sections. All protocols were performed in accordance with the Cell DIVE Platform (GE Research, Niskayuna, NY, USA). Slides were post-fixed using an ethanol-acetone (1:1) solution at 4°C for 1 minute. Slides were blocked overnight at 4°C in PBS with 3% BSA and 10% donkey serum (Bio-Rad), stained with DAPI (ThermoFisher) and mounted using mounting media (4% propyl gallate, 50% glycerol; Sigma-Aldrich).

Slides were imaged at 20X to acquire background autofluorescence, which was subtracted from the following staining round. Slides were decoverslipped in PBS; an antibody mixture (unconjugated primary antibodies for the first round of staining and conjugated antibodies for subsequent staining) was prepared and incubated for 1 hour at RT or overnight at 4°C. Slides were washed thrice in PBS for 5 minutes with gentle agitation. An antibody mixture with appropriate secondary antibodies was prepared and slides were incubated for 1 hour at RT, washed thrice with PBS, re-coverslipped, and imaged. After imaging, a bleaching round was performed by decoverslipping and incubating the tissue slides thrice for 15 minutes in 0.5M NaHCO_3_ (pH 11.2) and 3% H_2_O_2_ with a 1-minute wash in between, followed by 3 washes in PBS and a 2-minute DAPI recharge. Slides were then re-coverslipped and a bleached image was acquired which was subtracted from the following staining round. Staining and bleaching rounds were repeated until completion of 4 staining rounds (Table 3). Images were analysed using QuPath (v0.3.2) and representative images are shown.

**Table 3.**
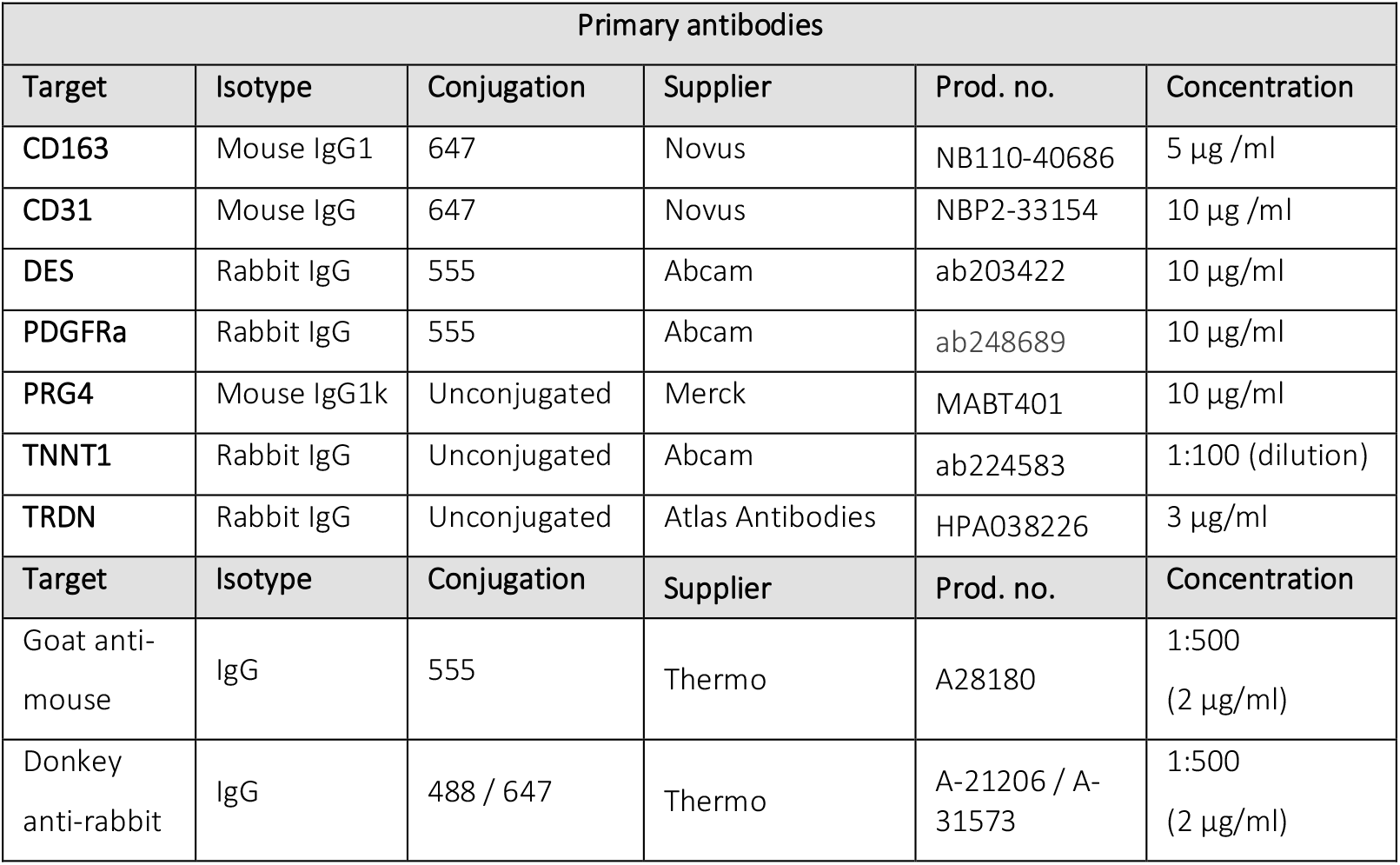
Overview of primary and secondary antibodies used for immunofluorescence staining.

### 2.6 Trichrome staining

Snap-frozen tendon samples were embedded in OCT before cutting 6 μm tissue sections. All steps were performed at room temperature. Slides were air-dried for 15 minutes, fixed in 10% formalin for 15 minutes and washed with ultrapure water. Nuclei were stained using Dako Mayer’s haematoxylin (Alignment Technologies) for 15 minutes, washed, blued with Dako Bluing Buffer (Alignment Technologies) and washed again. The Trichrome Stain Kit – Masson (RHS-773-LG) was used. Slides were stained with 1:1 Trichrome stain (A) and distilled water for 30 seconds, washed, and then treated with fresh Phosphotungstic/Phosphomolybdic Acid solution for 8 minutes (1 Phosphotungstic Acid: (B):1 Phosophomolybdic Acid (C): 2 distilled water). The slides were washed, then counterstained in 1: 1 Light Green (D) and distilled water for 10 seconds, and washed again. Slides were immersed in fresh 1% acetic acid solution for 8 minutes, washed, dehydrated, and mounted in DPX. Slides were imaged using the Motic EasyScan 1 system.

### 2.7 DAB staining

Paraffin-embedded hamstring was sectioned at 6 μm and dewaxed sections subjected to antigen retrieval using Vector Laboratories Tris-based (pH9.0) solution at 95°C for 20min. Sections were rested as the buffer cooled for 20min, blocked for endogenous peroxidase with 0.3%H2O2 for 10 minutes, and permeabilised with 0.5% Triton-X for 10 minutes. Non-specific binding was blocked for 1hr at RT using 3% Horse serum and primary antibody applied overnight at 4°C in a humidified chamber. Primary Pax 7 (ab218472) antibody was used at 1:1000 (0.2μg/ml) diluted in 3% BSA/1X PBS and 3% horse serum. Detection was carried out using the VECTASTAIN® Elite® ABC-HRP Kit with ImmPACT® DAB Substrate Kit and Vector Haematoxylin QS as counterstain. Slides were then dehydrated and mounted in DPX. Slides were imaged using the Motic EasyScan 1 system.

## 3 Results

### A single-nucleus transcriptomic atlas of healthy human hamstring tendon

To gain a comprehensive view of healthy human hamstring tendon, we utilised snRNA-seq and spatial transcriptomics, in combination with immunohistochemistry and immunofluorescence. Using sn-RNASeq, we profiled the transcriptomes of 10,533 nuclei from 4 healthy donors. Isolated nuclei from each sample were subjected to droplet-based 3’ end sequencing (10X Chromium). After batch correction and integration using Harmony (Supplementary Figure 3), we performed unsupervised graph clustering and annotation of our individual cell populations (Figure 1A), each displaying canonical marker genes (Figure 1B). We identified a range of different cell types, including 2 fibroblast subsets (*COL1A1, COL1A2, COL3A1, DCN*), 3 skeletal muscle cell clusters (*TRDN, DES*), satellite cells (*PAX7, CALCR, GREM1*), 2 endothelial clusters (*PECAM1, PTPRB, VWF*), mural cells (*NOTCH3, PDGFRB, MYO1B*), adipocytes (*GRAM*, AQP7, *ADIPOQ, PLIN1*), immune cells (*PTPRC, CD247, CD69, BLNK, CD163*), and “nerve cells”, which are undefined nervous system cells, expressing markers including *TENM2, NTRK3, NRP2, COL23A1, ROR1*, and *GRID2*. The two fibroblast subsets are subsequently referred to as MKX+ and PDGFRA+ fibroblasts, due to their expression of these markers (see also Figure 3). Fast- and slow-twitch skeletal muscle cells were identified, along with a third cluster which likely represents cells transitioning between these two states (see also Figure 4). All cell populations were represented across all healthy donors, apart from the slow-twitch skeletal muscle cells and adipocytes, which were only present in 3/4 donors (Supplementary Figure 5).

**Figure 1.**
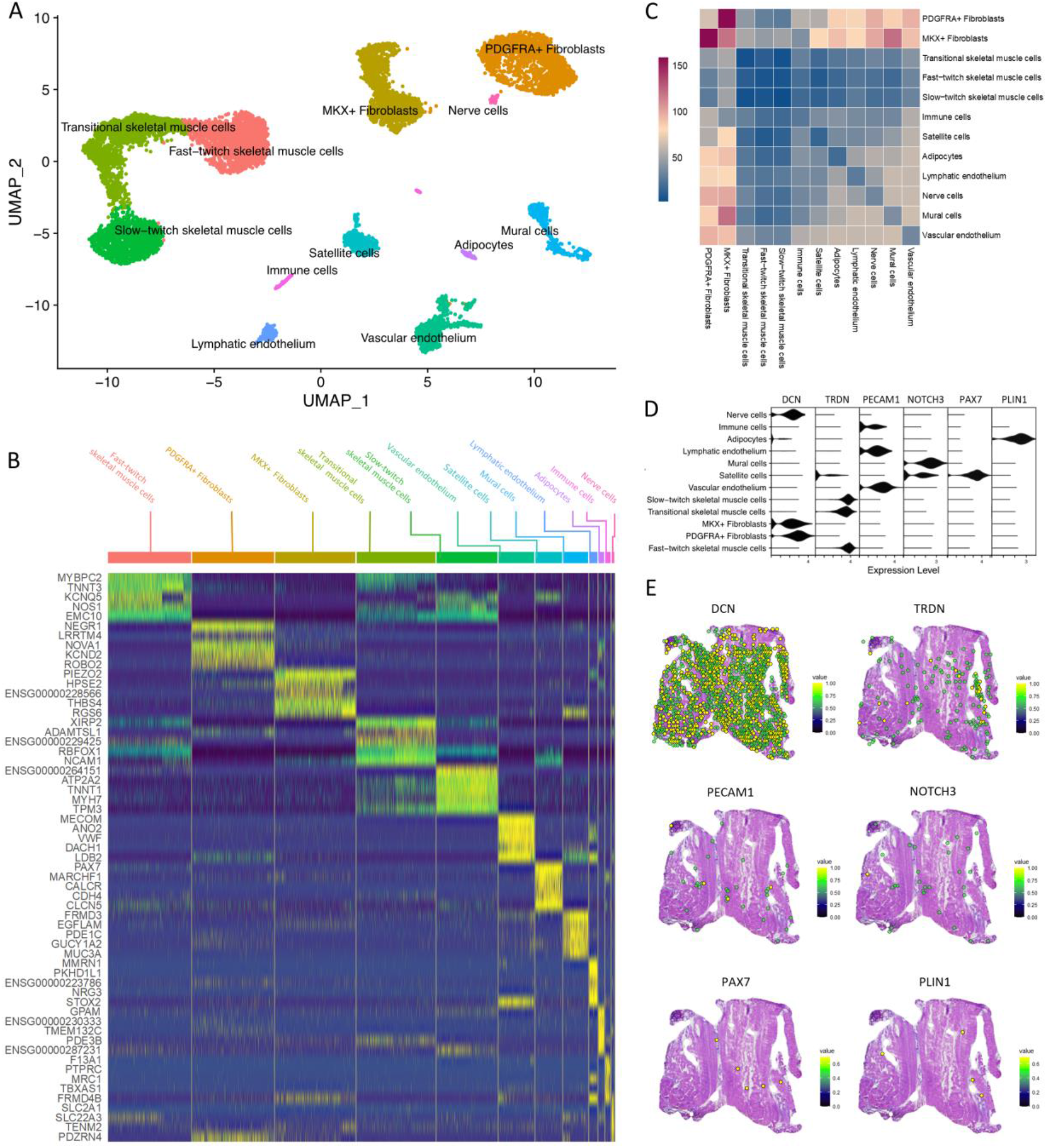
Transcriptomic analysis reveals 12 different cell types in healthy hamstring tendon. (A) Uniform manifold approximation and projection (UMAP) embedding of 10,533 cells from 4 individuals reveals 12 clusters. (B) Heatmap of the top 5 positive differentially expressed genes for each cluster. If gene symbol names were not available, ensemble names were used. (C) CellphoneDB analysis of predicted ligand-receptor interactions between different cell types in the snRNA-seq datasets. (D) Expression of *DCN* (fibroblasts), *TRDN* (skeletal muscle cells), *PECAM1* (endothelial cells), *NOTCH3* (mural cells), *PAX7* (satellite cells), and *PLIN1* (adipocytes) in snRNA-seq datasets. (E) Spatial expression of *DCN, TRDN, PECAM1, NOTCH3, PAX7*, and *PLIN1* in the healthy hamstring.

CellPhoneDB analysis was used to predict the potential ligand-receptor interactions within and between the identified cell types. This analysis revealed that the two fibroblast subsets had the highest number of ligand-receptor interactions (165), showing interactions within and between each fibroblast cluster, and with other clusters such as adipocytes, satellite cells, and endothelial cell subsets, suggesting that fibroblasts are key regulators of hamstring tendon tissue homeostasis (Figure 1C). All three skeletal muscle cell types and immune cells have very few predicted interactions with the same or other clusters. All other cell types have some predicted interactions with each other, but have the highest number of potential interactions with the two fibroblast cell types.

To acquire an understanding of the spatial distribution of cells across the hamstring tendon tissue we performed spatial transcriptomics (10X Visium, n=1). Canonical gene expression markers that were used to identify the major cell types in the snRNA-seq data (*DCN* for fibroblasts, *TRDN* for skeletal muscle cells, *PECAM1* for endothelial cells, *NOTCH3* for mural cells, *PAX7* for satellite cells, and *PLIN1* for adipocytes) (Figure 1D) were also identified in the spatial transcriptomics data (Figure 1E). Overall, our snRNA-seq datasets identify major cell types residing within healthy human hamstring tendon, including skeletal muscle cells, adipocytes, and nerve cells, which have not been identified with transcriptomic methods previously.

### Mechanistic insights into the regulatory networks driving healthy hamstring tendon cellularity

We next sought to evaluate gene regulatory network reconstruction and cell-state identification using SCENIC. SCENIC evaluates which regulons, or collections of transcription factors and co-factors, are significantly enriched in each cell. We identified 317 regulons with significantly enriched motifs, and the resulting binary regulon activity matrix (Figure 2A) clearly stratifies similar cell types identified by clustering the integrated snRNA-seq data. The three skeletal muscle cell clusters, located on the left side of the matrix, have the most distinct regulon activity when compared to all the other clusters. Interestingly, satellite cells, multipotent cells present in skeletal muscle, did not cluster with the skeletal muscle cells. Both of the fibroblast cell types had distinct sets of activated regulons. Cell clustering based on regulon activity reflected the findings from the regulon activity matrix (Figure 2B). All cell subsets, apart from the small nerve cells cluster, could easily be detected as a separate cluster. The three skeletal muscle cell types, especially the slow-twitch and fast-twitch skeletal muscle cell subsets, showed a lot of overlapping regulon activity but were different from non-skeletal muscle cells. Satellite cells clustered close to the fibroblast subsets rather than close to the skeletal muscle cell subsets. Although the two fibroblasts cell clusters were present near each other, they were clearly distinct. Finally, the regulon specificity score (RSS) was calculated to assess regulon-cell type cluster specificity, and the top 5 regulons per cell type were highlighted (Supplementary Figure 6).

**Figure 2.**
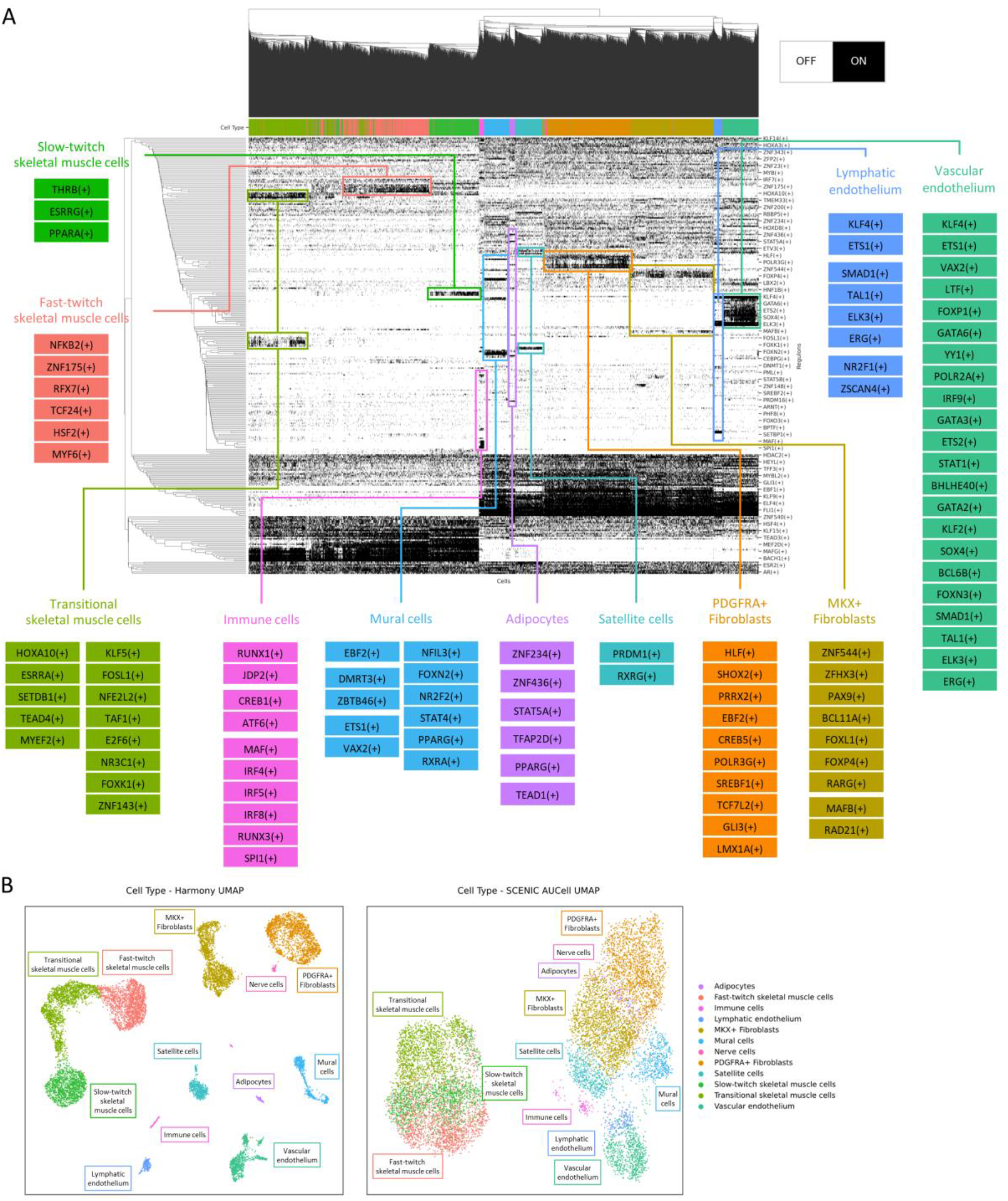
SCENIC analysis provides mechanistic insights into the regulatory networks driving healthy hamstring tendon function (A) Binary regulon activity matrix of the 317 regulons that were found to have significantly enriched motifs for the corresponding transcriptions factors. (B) Comparison of the UMAP using Harmony integration (left) or the SCENIC regulon activity matrix (right).

### Two distinct fibroblast cell types are detected in healthy hamstring tendon

In order to better understand the similarities and differences between the MKX+ and PDGFRA+ fibroblasts, we compared their gene expression profiles (Figure 3A). Markers previously described as specific for tendon fibroblasts, including mohawk (*MKX*), tenomodulin (*TNMD*), thrombospondin-4 (*THBS4*), and cartilage oligometric matrix protein (*COMP*), were all exclusively present in MKX+ fibroblasts. Other highly expressed genes included piezo-type mechanosensitive ion channel component 2 (*PIEZO2*), which is important for proprioception, and different collagens and ECM proteins (*COL11A1, COL11A2, COL12A1, COL14A1, FMOD*). In contrast, PDGFRA+ fibroblasts specifically express high levels of neuronal growth regulator 1 (*NEGR1*), RNA-binding protein Nova-1 (*NOVA1*), and vitrin, as well as fibrillin-1 (*FBN1*), fibulins (*FBLN1, FBLN2, FBLN5*), and elastin (*ELN*), which are all key components of elastic fibres. Differences between these two fibroblasts subsets are further illustrated by the differential expression analysis (Figure 3B). Pathway analysis using highly expressed genes for each fibroblast subset revealed that both cell types had strong upregulation of “ECM organisation”, “collagen formation”, “collagen biosynthesis and modifying enzymes”, and “ECM proteoglycans”. However, increased expression of “elastin fibre formation” and “molecules associated with elastic fibres” were only found in PDGFRA+ fibroblasts, while “signalling by receptor tyrosine kinases”,” signalling by MET”, “MET promotes cell motility”, and “MET activates PTK2 signalling” were only significant in MKX+ fibroblasts (Supplementary Figure 7). Immunofluorescence staining of healthy human hamstring tendon showed that these tendon fibroblasts, recognisable by their typical elongated shape, are positive for fibroblast marker lubricin (PRG4), while only a subset of fibroblasts showed expression of PDGFRA (Figure 3C). We performed cell2location analysis to leverage the cell type annotations identified in the snRNA-seq data and decompose the cell type spatial location of each cluster. We found that both fibroblast cell types, but in this tissue mainly MKX+ fibroblasts, are predicted to be located throughout the whole hamstring, while the PDGFRA+ fibroblasts were predicted to be especially abundant in areas close to skeletal muscle (Figure 3D). Although tendon fibroblasts are thought to be mainly responsible for the production and organisation of ECM in tendon, not only fibroblasts, but other cell types such as satellite cells, mural cells, adipocytes, and nerve-like cells were also found to be potential producers of type I and type III collagens (Figure 3E). Pathway analysis confirmed that while both fibroblast subsets had a high enrichment score for ECM-related GO:BP terms, other cells, including satellite cells, vascular and lymphatic endothelium, mural cells, and nerve-like cell, are also involved in these processes (Figure 3F).

**Figure 3.**
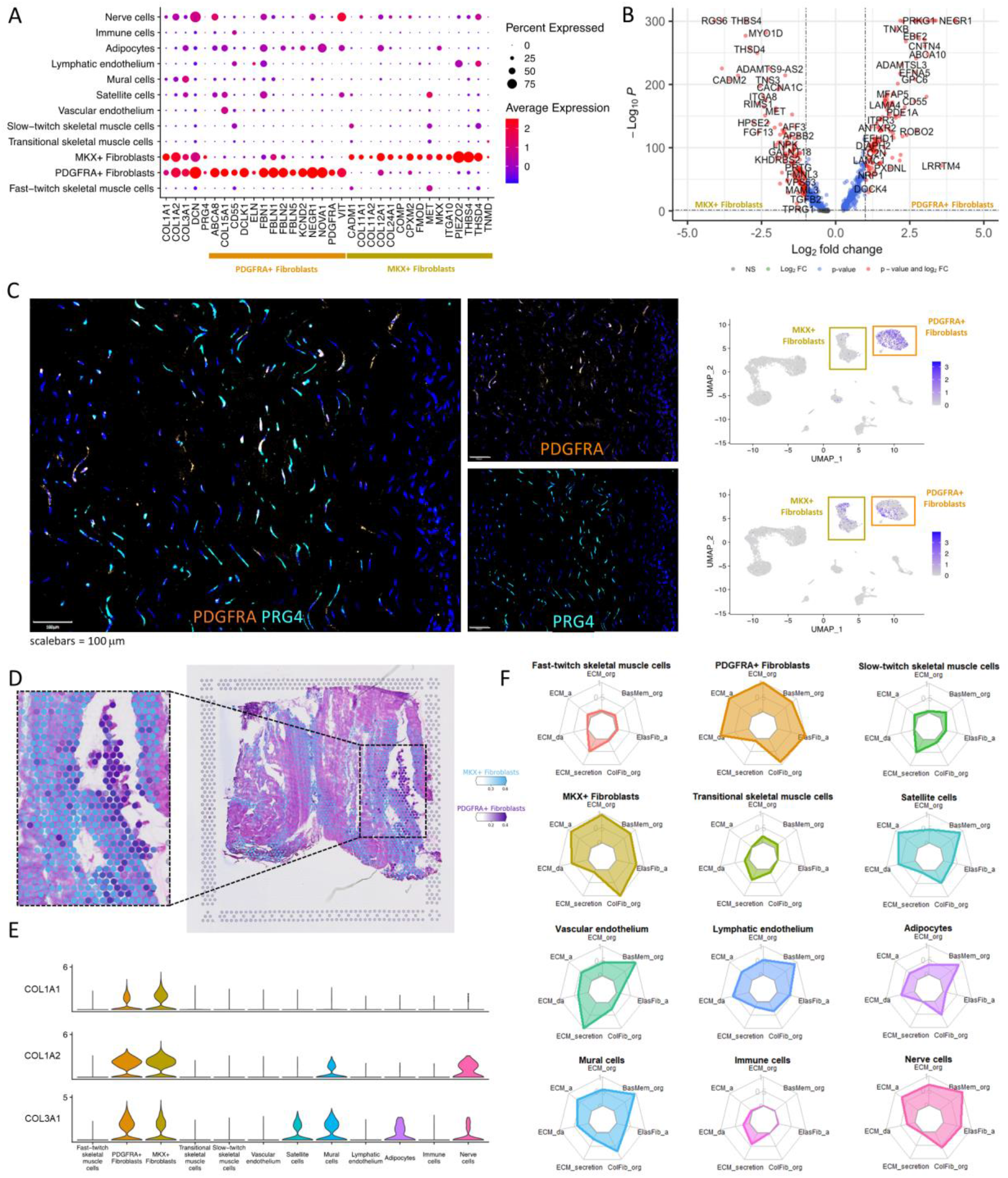
Healthy hamstring tendon contains two different fibroblast types. (A) Expression of fibroblast markers across all different cell types in the snRNA-Seq datasets. (B) Differential expression analysis of PDGFRA+ Fibroblasts and MKX+ Fibroblasts, grey = not significant, green = log2 fold change (Log2FC) at least ±1, but p > 0.05, blue = p < 0.05 but Log2FC between 1 and –1, red = p < 0.05 and Log2FC at least ±1. (C) Expression of fibroblast marker lubricin (PRG4) and PDGFRA+ fibroblasts marker platelet derived growth factor receptor alpha (PDGFRA) in healthy human tendon tissue compared to their RNA expression in the feature plots. (D) Predicted location of MKX+ fibroblasts (blue) and PDGFRA+ fibroblasts (purple) in the spatial transcriptomics data (Visium) using cell2location analysis. (E) Expression *COL1A1, COL1A2*, and *COL3A1* in snRNA-seq datasets. (F) Radar plots of the enrichment score (0-1) of gene ontology biological processes (GO:BP) related to extracellular matrix (ECM). Clockwise starting at the top: ECM organisation, basement membrane organisation, elastic fibre assembly, collagen fibre organisation, ECM secretion, ECM disassembly, ECM assembly.

### Three types of skeletal muscle-like cells are present in human hamstring tendon

Human tendons are highly transitional tissues that transition from bone attachment (enthesis) through tendon midbody to the muscle attachment (myotendinous junction, MTJ). The tissue used in this study includes the region of hamstring tendon from the midbody to the MTJ. Although macroscopic muscle at the MTJ was removed from tendon samples during tissue processing for sequencing, tendon regions are a continuum and it is unsurprising that skeletal muscle was present in this area. Three cell types were identified with high expression of the skeletal muscle markers triadin (*TRDN*) and desmin (*DES*)(Figure 4A-C). Two of the three clusters expressed high levels of troponins: while one cluster expressed high levels troponin isoforms associated with fast-twitch skeletal muscle (*TNNT3, TNNC2*, and *TNNI2*), the other cluster expressed high levels of isoforms associated with slow-twitch skeletal muscle (*TNNT1, TNNI1*, and *TNNC1*). The third cell type was not associated with high expression of troponin isoforms, but instead expressed high levels of *COL22A1*, a transcript for an ECM protein which is mainly located in tissue junctions. Therefore, these cells were designated “transitional skeletal muscle cells”. Trichrome staining clearly shows the presence of skeletal muscle (red) infiltrating the collagen dense matrix of tendon (Figure 4D). Furthermore, immunofluorescence confirms the presence of both triadin (TRDN) and troponin T1 (TNNT1) in healthy hamstring tissue (Figure 4E). Visually, all three skeletal muscle cell types are predicted to be located simultaneously within one region (as highlighted). In addition, each skeletal muscle cell type is also located throughout some other parts of the tissue section (Figure 4F). Pathway analysis revealed that both fast- and slow-twitch skeletal muscle cells were associated with “muscle contraction”, “striated muscle contraction”, and “ion homeostasis”, while no Reactome pathways were significantly enriched in the transitional skeletal muscle cells (Figure 4G). As all three of these cell types only have a limited number of potential ligand-receptor interactions (Figure 1C), it is more likely that their interactions are mechanical interactions rather than molecular in hamstring tendon MTJ.

**Figure 4.**
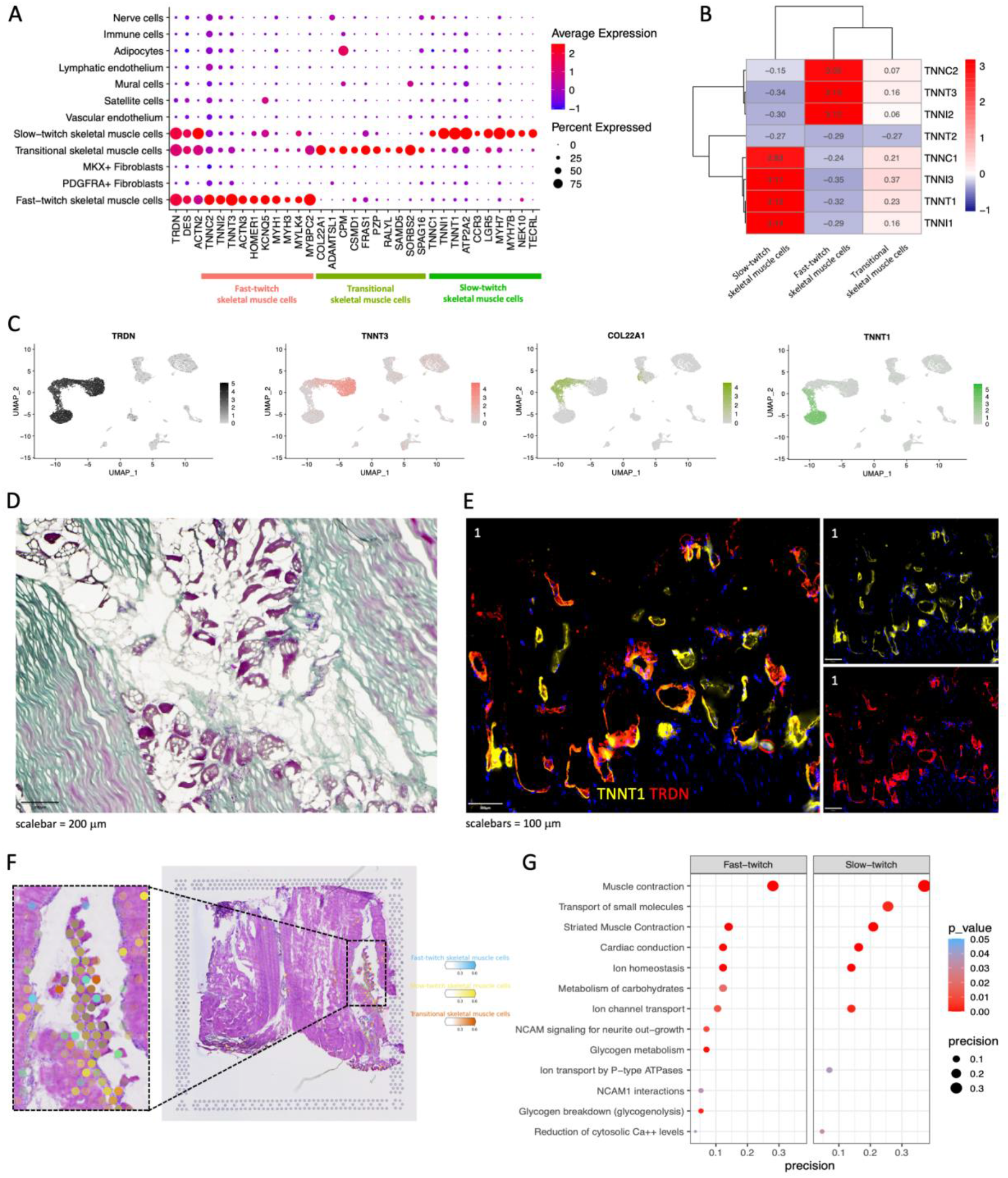
Three types of skeletal muscle cells can be identified in healthy human hamstring tendon using snRNA-seq. (A) Highly expressed genes across the three types of skeletal muscle cells. (B) Heatmap of the expression of all troponins across the three skeletal muscle cell types, identifying fast- and slow-twitch skeletal muscle cells. (C) Feature plots of the expression of *TRDN, TNNT3, COL22A1*, and *TNNT1* in the snRNA-seq datasets. (D) Trichrome staining of human healthy hamstring tendon tissue, including muscle (red) and collagen (green). (E) Immunofluorescence staining of troponin T1 (TNNT1, yellow) and triadin (TRDN, red) in human healthy hamstring tendon tissue. (F) Cell2location prediction of the location of the fast-twitch skeletal muscle cells (blue), slow-twitch skeletal muscle cells (yellow), and transitional skeletal muscle cells (orange) using the spatial transcriptomics (10X Visium) data. (G) Enriched pathways in the Reactome database for each of the three skeletal muscle cell types (no significant pathways for transitional skeletal muscle cells).

### The vascular compartment of hamstring tendon is composed of three distinct populations

The vascular compartment includes 3 distinct populations: vascular endothelium (*FLT1, CD34, NOTCH4*), lymphatic endothelial (*MMRN1, PROX1, KDR, FLT4*), and mural cells (*NOTCH3, PDGFRB, MYO1B*) (Figure 5A). Cell2location analysis demonstrates a large overlap in the predicted locations of vascular endothelial cells (black), mural cells (blue), and lymphatic endothelium (orange), suggesting possible interactions between these cell types (Figure 5B, Figure 1C). Immunofluorescence confirms the presence of vessels across healthy hamstring tendon tissue. However, vessels seem to be much more abundant in areas close to muscle fibres than in the main body of the tendon (Figure 5C). GO:BP pathways associated with vascular processes, including cell migration, cell communication, signalling, blood vessel development, angiogenesis, chemotaxis, and cell motility, were all strongly enriched in vascular endothelium. Strong enrichment of most of these pathways was also found in all other cell types, except for skeletal muscle cells and satellite cells, which only showed minimal enrichment (Figure 5D). Enriched pathways from the Reactome database were “signal transduction” and “signalling by Rho GTPases” for vascular endothelium, “signal transduction” and “signalling by GPCR” in mural cells, and “RHOG GTPase cycle” in lymphatic endothelium (Supplementary Figure 8). The predicted ligand-receptor interactions show a high number of predicted interactions between the three vascular cell types and the two fibroblast cell types, especially between mural cells and MKX+ fibroblast (Figure 1D). Among the top predicted ligand-receptor interactions from mural cells to MKX+ fibroblast are pairs involved in the GO:BP pathways signal transduction (COPA & EGFR), angiogenesis (FGF1 & TGFBR3) and BMP signalling (BMP5 & multiple receptor complexes). Among the top predicted ligand-receptor interactions from MKX+ fibroblasts to mural cells are immune response / cell adhesion (THBS1 & CD36), FGF2 & CD44 which are involved in ECM organisation and regulation of ERK pathways, and BMP signalling (BMP5 and BMP6 & multiple receptor complexes) (Supplementary Table 1).

**Figure 5.**
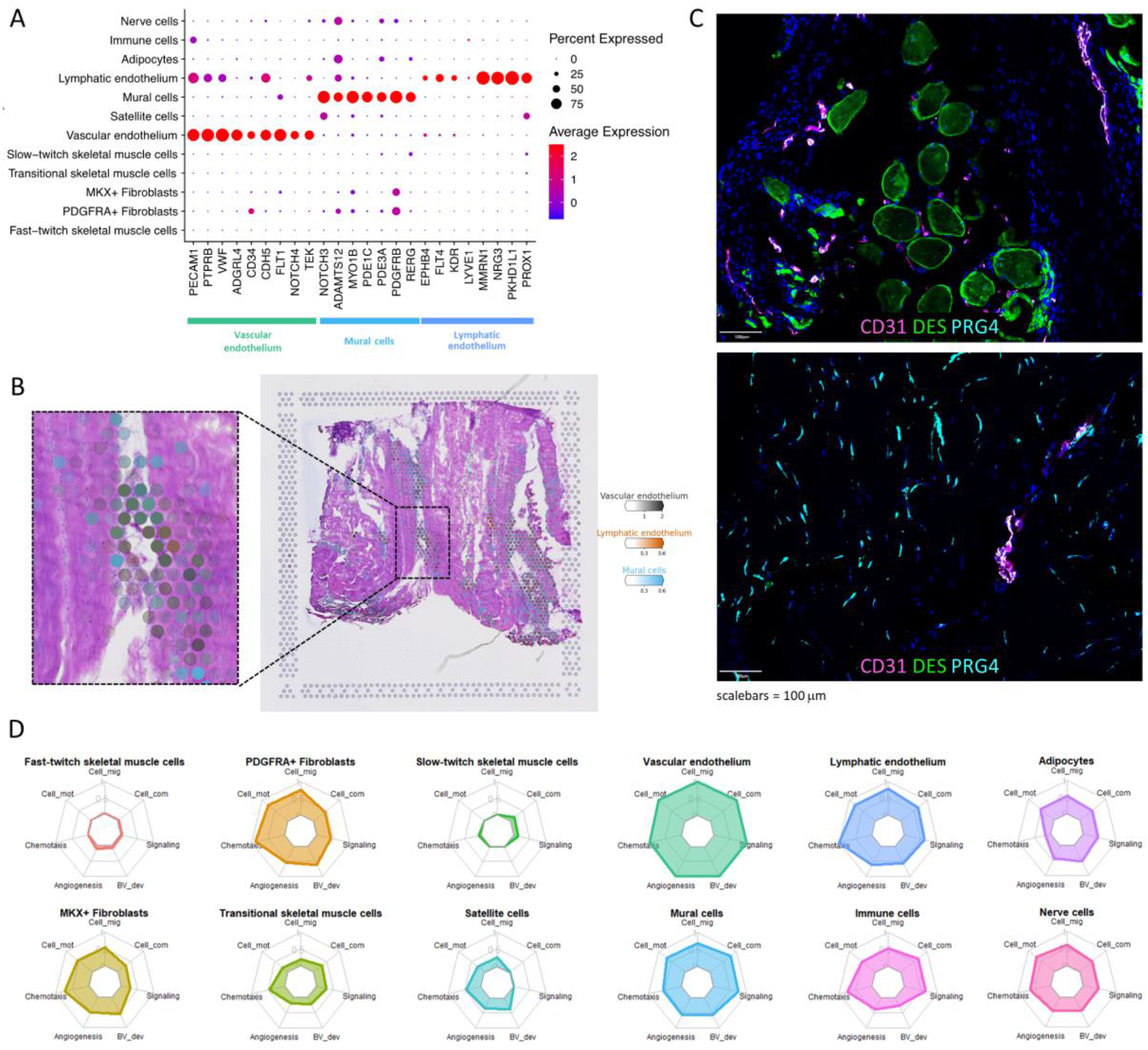
Vascular compartment in healthy hamstring tendon. (A) Expression of endothelial and mural cell markers across all different cell types in the snRNA-Seq datasets. (B) Predicted location of vascular endothelium (black), lymphatic endothelium (orange), and mural cells (blue) fibroblasts in the spatial transcriptomics data (Visium) using cell2location analysis. (C) Immunofluorescence. (D) Radar plots of the enrichment score (0-1) of gene ontology biological processes (GO:BP) related to vascular function. Clockwise starting at the top: cell migration, cell communication, signalling, blood vessel development, angiogenesis, chemotaxis, and cell motility.

### Macrophage, T cell, and dendritic cells are present in healthy hamstring tendon

Re-clustering of the identified immune cells (108 nuclei, ^~^1% of all nuclei) revealed two distinct cell clusters representing three different cell types. The three immune cell subsets were identified as macrophages (*CD14*, *CD163, MERTK, MRC1, MSR1*), T cells (*CD247, IL7R, CD69, THEMIS*), and dendritic cells (*HLA-DQA1, CD74, GPAT3, CIITA, HLA-DRB1*) (Figure 6A-C). Although further delineation of the T cell clusters was not possible due to the small number of cells and the low expression of CD4 and CD8 classical T cell markers, the expression of *CD69* could indicate that at least a subset of these cells are tissue-resident memory T cells. The strong expression of *MERTK* in the macrophage population also suggest that this population is mostly comprised of tissue-resident macrophages. Cell2location predicts the presence of immune cells in similar areas as vascular endothelium, but also in areas where skeletal muscle cells are predicted to be located (Figure 6D). Presence of immune cells in hamstring tissue was confirmed by immunofluorescence staining, which showed that immune cells could mainly be found around vessels and colocalization of CD163 macrophages with CD3 T-cells (Figure 6E). A relatively low number of potential ligand-receptor interactions were identified for immune cells, indicating a low activation state of the immune cells in our datasets, consistent with the samples coming from healthy hamstring (Figure 6F). Reactome pathways enriched in immune cells included “Signal Transduction”, “Innate Immune System”, and “Adaptive Immune System” (Supplementary Figure 9).

**Figure 6.**
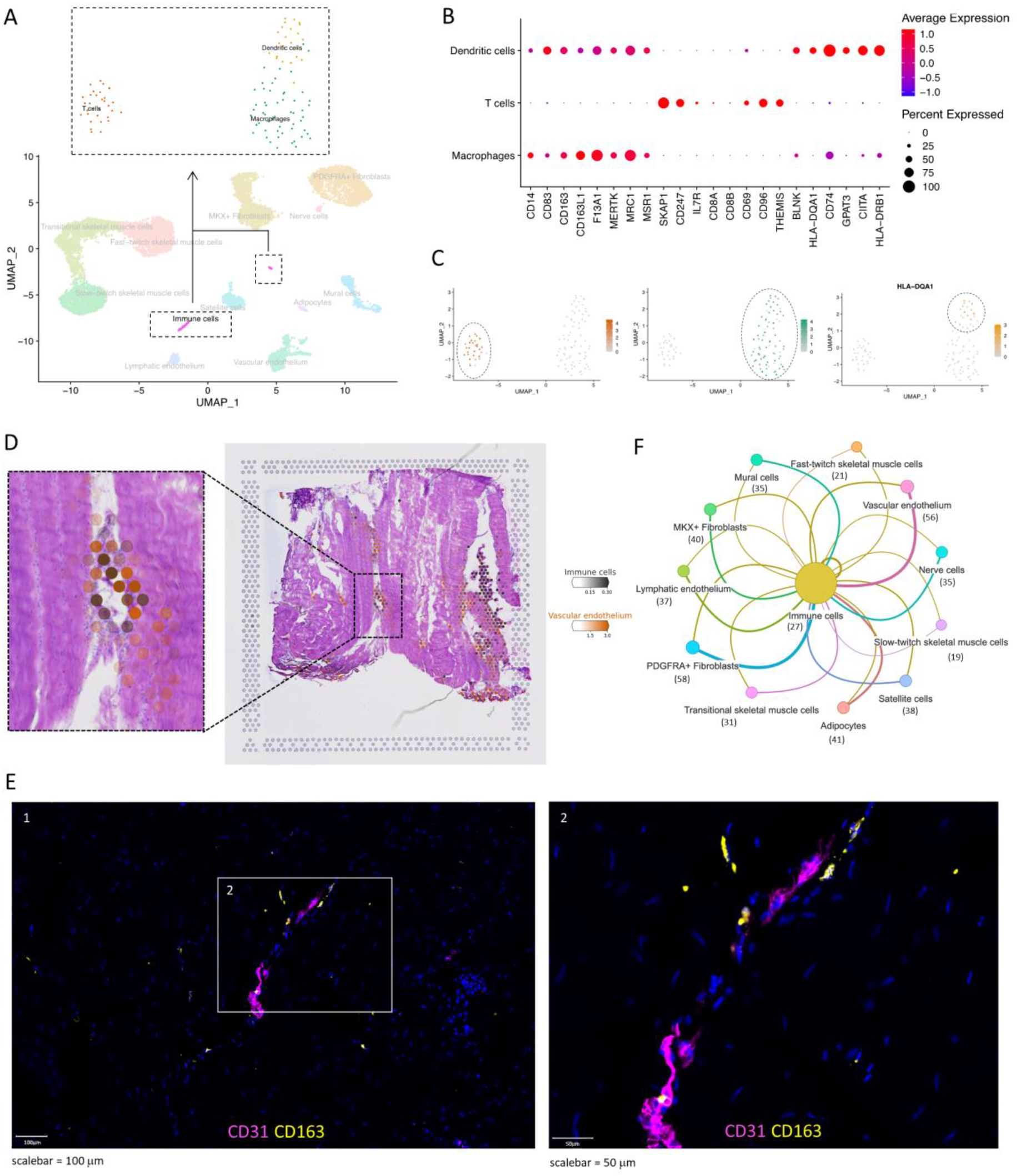
Immune cells in healthy hamstring tendon. (A) The immune cell cluster was re-clustered, revealing two distinct clusters representing T cells and macrophages. (B) Expression of immune cell markers in the two cell types within the immune cell cluster. (C) Featureplots of *SKAP1* and *MRC1* in the immune cell cluster. (D) Cell2location prediction of the location of immune cells and vascular endothelium using the spatial transcriptomics (10X Visium) data. (E) Immunofluorescence confirms the presence of CD163+ macrophages in healthy hamstring tendon tissue. (F) Predicted number of interactions of immune cells with the other cell types within the snRNA-seq datasets using CellPhoneDB and InterCellar.

### Adipocytes, satellite cells, and nerve cells are found close to fibroblasts in healthy hamstring tendon

Finally, three cell types that were not previously reported in human tendon single cell transcriptomics studies, namely adipocytes, satellite cells, and nerve cells, which are undefined nervous system cells, were identified in our datasets (Figure 7A). This small cluster of nerve cells, or undefined nervous system cells, represented only ^~^0.5% of all cells and expressed markers related to both nerve cells and glial cells. Adipocytes and satellite cells were identified by expression of known markers (Figure 7A). Cell2location analysis (Figure 7B) showed that adipocytes were predicted to be located in similar areas as PDGFRA+ fibroblasts and skeletal muscle cells (as per Figure 3D and Figure 4F, respectively), and satellite cells and nerve cells were predicted to be located in similar areas as fibroblasts and cells from the vascular compartment (as per Figure 3D and 5B, respectively). Immunohistochemistry staining of PAX7 revealed that satellite cells are present in close proximity to muscle fibres within tendon tissue (Figure 7C). No enriched Reactome pathways were found for satellite cells. However, Reactome pathways “Metabolism” and “Metabolism of lipids” were significantly enriched in adipocytes and “Signal Transduction” in nerve cells, among others (Figure 7D).

**Figure 7.**
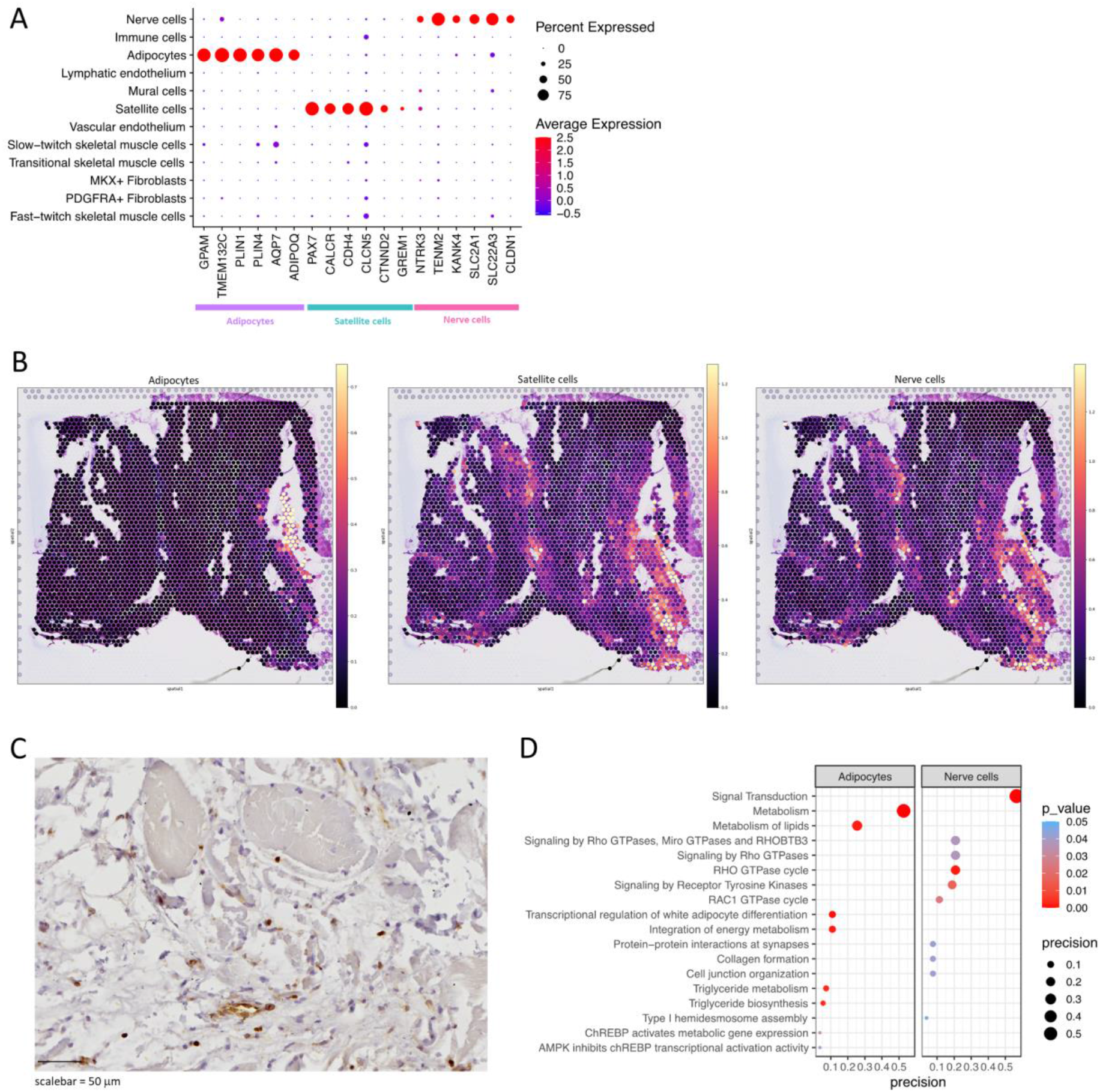
Adipocytes, satellite cells, and nerve cells in healthy human hamstring tendon. (A) Expression of highly expressed markers in adipocytes, satellite cells, and nerve cells. (B) Cell2location analysis predicted location of adipocytes, satellite cells, and nerve cells. (C) Reactome results of enriched pathways for adipocytes and nerve cells (no significant pathways for satellite cells). (D) Immunohistochemistry staining for PAX7 reveals that satellite cells are present in close proximity to muscle fibres in tendon tissue.

## Discussion

In this study, we systematically characterised healthy human hamstring tendon using snRNA-seq, spatial transcriptomics, and imaging, resulting in a comprehensive atlas that can act as a reference map for future studies. SnRNA-seq was utilised in this study as it can deliver considerably enhanced extraction of nuclei from dense collagen-rich tissue and the removal of a stress-response to prolonged enzymatic digestion. Unsupervised clustering was used to identify 12 cell types, which were annotated according to known markers from the literature, differentially expressed genes, and pathway analysis. We confirmed the presence of two distinct fibroblast subsets along with endothelial cells, mural cells, and immune cells, and revealed the presence of less well-characterised cells, including different skeletal muscle cell types, satellite cells, adipocytes, and nerve cells. The low frequency of immune cells in our datasets (^~^1%) corroborates that this is indeed healthy tissue. Our datasets suggest that fibroblasts are key regulators of hamstring tendon tissue homeostasis due to their important role in the production and organisation of ECM as well as due the high number of potential ligand-receptor interactions, both within each fibroblast cell type as well as with all other cell types.

Our datasets show both similarities and differences with previous human single cell transcriptomics studies[8, 9]. These studies identified several different fibroblasts, mural cells, endothelial cells, and immune cells, including monocytes/macrophages, T cells, and dendritic cells. Most markers that were used to differentiate between different tenocyte populations in the paper by Kendal *et al*. were present in either both or neither of our fibroblast populations; markers used by Akbar *et al*. to identify normal tenocyte1 and normal tenocyte2 mostly aligned with our PDGFRA+ fibroblasts and MKX+ fibroblasts, respectively (Supplementary Figure 10). However, these studies did not detect the presence of adipocytes and nerve cells in their tissues and have low representation of skeletal muscle cells or satellite cells. Differences between this study and previous studies could have arisen due to the fact that a greater number of healthy nuclei was analysed in our study or due to the differences in transcriptomics methods (single cell vs single nucleus approaches). While single cell transcriptomics generally capture more RNA per cell, including stable transcripts in the cytoplasm that might not be present in the nucleus, single nucleus transcriptomics enables the capture of a wider variety of cells, including cells that are hard to capture with single cell techniques due to the buoyancy or larger size of cells[26, 27].

Due to the origin of the tissue, it is unsurprising that skeletal muscle cells were found in hamstring tendon samples. The multinucleated nature of skeletal muscle combined with the low cellularity of tendon could explain the relatively high percentage of skeletal muscle cells (^~^45%) in our datasets. The limited number of predicted ligand-receptor interactions of skeletal muscle cell types with all other cell types suggests that skeletal muscle fibres might be present in healthy hamstring MTJ mostly for mechanical attachment. It is interesting to note that the PDGFRA+ fibroblast subset enriched in elastin biology sits spatially closer to the muscle and further study would be warranted to establish functional specialisation linked to mechanical attachment. The single cell transcriptomics study by Akbar *et al*. also identified a small muscle subset in their tendon datasets. However, the three markers defining the muscle cell subset were all present in the satellite cells but not skeletal muscle cells in our dataset, suggesting that these might have been satellite cells[9]. The satellite cells in our datasets did not cluster together with skeletal muscle cells and their regulon activation profile was more similar to non-skeletal muscle cells such as mural cells, endothelial cells, and fibroblasts, than skeletal muscle cells. Interestingly, a previous study has shown that there is a loss of satellite cells in skeletal muscle after tendon injury[28], illustrating that these cells can respond to changes in tendon. However, more work is needed to better understand the role and localisation of satellite cells as well as the potential interactions of these presumably skeletal muscle fibre-based cells with tendon-based cells.

Analysis of regulatory networks using SCENIC corroborated our snRNA-seq cell type definitions and also gave further insight into the potential functions of each identified cell type, especially the two fibroblast subsets. Regulons upregulated in the MKX+ fibroblasts include PAX9, which is involved in tendon development [29]. However, most other regulons have previously been associated with other tissues: for example MAFB, which regulates macrophage differentiation[30], and ZFHX3, which is associated with neurons[31]. The link with neurons is intriguing as these MKX+ fibroblasts expressed high levels of *PIEZO2*, piezo-type mechanosensitive ion channel component 2, which is known to be critical for proprioception and has previously been identified in the Golgi tendon organs in mice[32]. Further studies will be required to validate the function of these regulons in tendon fibroblasts. In the PDGFRA+ fibroblasts, the activated regulons include SHOX2, PRRX2, EBF2, CREB5, and TCF7L2, which are known to be involved in musculoskeletal and (early) limb development, as well as tendon healing; many of these factors are also involved in or can affect wnt/b-catenin signalling [33–39]. Interestingly, PDGFRA+ cells have previously been identified as markers for progenitor cells in human skeletal muscle[40, 41], and *Tppp3+ Pdgfra+* tendon cells have previously been identified as tendon-derived stem cells (TDSC) in mice[42]. While only a small percentage of the PDGFRA+ fibroblasts in our datasets expressed *TPPP3*, this PDGFRA+ fibroblast population warrants further investigation in larger datasets to explore the potential progenitor-like function of (a subpopulation) of these cells.

Limitations of this study include the small number of included samples in transcriptomic methods. Although different cell types were identified that have not been reported on earlier, the addition of further samples would enable annotation of cell types with more granularity, as well as comparisons of features such as donor sex, age, and side of tendon. The possible functions of and interaction between different cell types were predicted based on mRNA expression only. Two difficulties with this are that snRNA-seq might not cover stable transcripts that are only present in the cytoplasm, and that the presence or lack of mRNA expression does not always translate to the presence or lack of protein expression. Therefore, additional studies are needed to validate cell-cell interactions and discover their functional effects. This might especially give more insight into the differences and similarities of the two different fibroblast cell types identified in this study. While spatial transcriptomics data was able to add some information about the presence and distribution of different cell types across the tissue, analysis is limited due to use of one sample in one tissue slice orientation. The lack of single-cell resolution in spatial transcriptomics and difficulty in accurate normalisation due to high variability in RNA expression levels between muscle and tendon areas were also limiting. The low cellularity of tendon and low transcriptional activity of the cells results in low detection of RNA in many parts of the tissue. The large differences in expression between high and low cellularity regions of the tissue could have biased the predicted locations of cells. Therefore, other spatial analysis techniques might be better suited to understand the spatial distribution of cells in tendon. Cell subsets were identified using RNA markers, and where possible these were validated by imaging methods. However, in some cases we were unable to obtain suitable antibodies (such as for mohawk).

In conclusion, this study enhances our understanding of cellular composition of healthy human hamstring tendon by using both transcriptional and spatial analyses. We show that healthy human hamstring tendon is comprised of wide range of cell types, including both previously demonstrated cell types such as fibroblasts, endothelial cells, mural cells, and immune cells, as well as less well-characterised cells, including different skeletal muscle cell types, satellite cells, adipocytes, and nerve cells. We identified two distinct types of fibroblasts, which are suggested to be the major regulators of tendon tissue homeostasis due their role in the production and organisation of ECM, the high number of predicted interactions with all other cell types, and the wide distribution across the tissue. Our datasets form the foundation of a comprehensive cellular atlas of healthy tendons that can act as a reference map for future studies, and will help dissect mechanisms of disease pathogenesis and identify new therapeutic targets.

## Data availability statement

All the code used for the data analysis of this paper is available here: https://github.com/Botnar-MSK-Atlas/hamstring_atlas. All the data in this manuscript will be openly available on Lattice on final publication and is hosted on Lattice. For access prior to final publication please contact the authors.

## Acknowledgements

We would like to thank our research assistant Louise Appleton, our research nurses Debra Beazley, Bridget Watkins, Kim Wheway, and Lois Vesty-Edwards, and the knee surgeon team at the Nuffield Orthopaedic Centre, for their invaluable help collecting human tissue for this study. We would like to thank the CellDive team at the Kennedy Institute of Rheumatology, including Mark Coles, Dylan Windell, and Ananya Bhalla, for their help with the immunofluorescence staining using the CellDive system. Our thanks also goes to Dr Carla Cohen for helpful discussions and for reading of the manuscript. Finally, we would like to thank all other members of the CZI Tendon Seed Network for insightful discussions.

