## Supplementary Figure for "Single nucleus and spatial transcriptomic profiling of human healthy hamstring tendon"

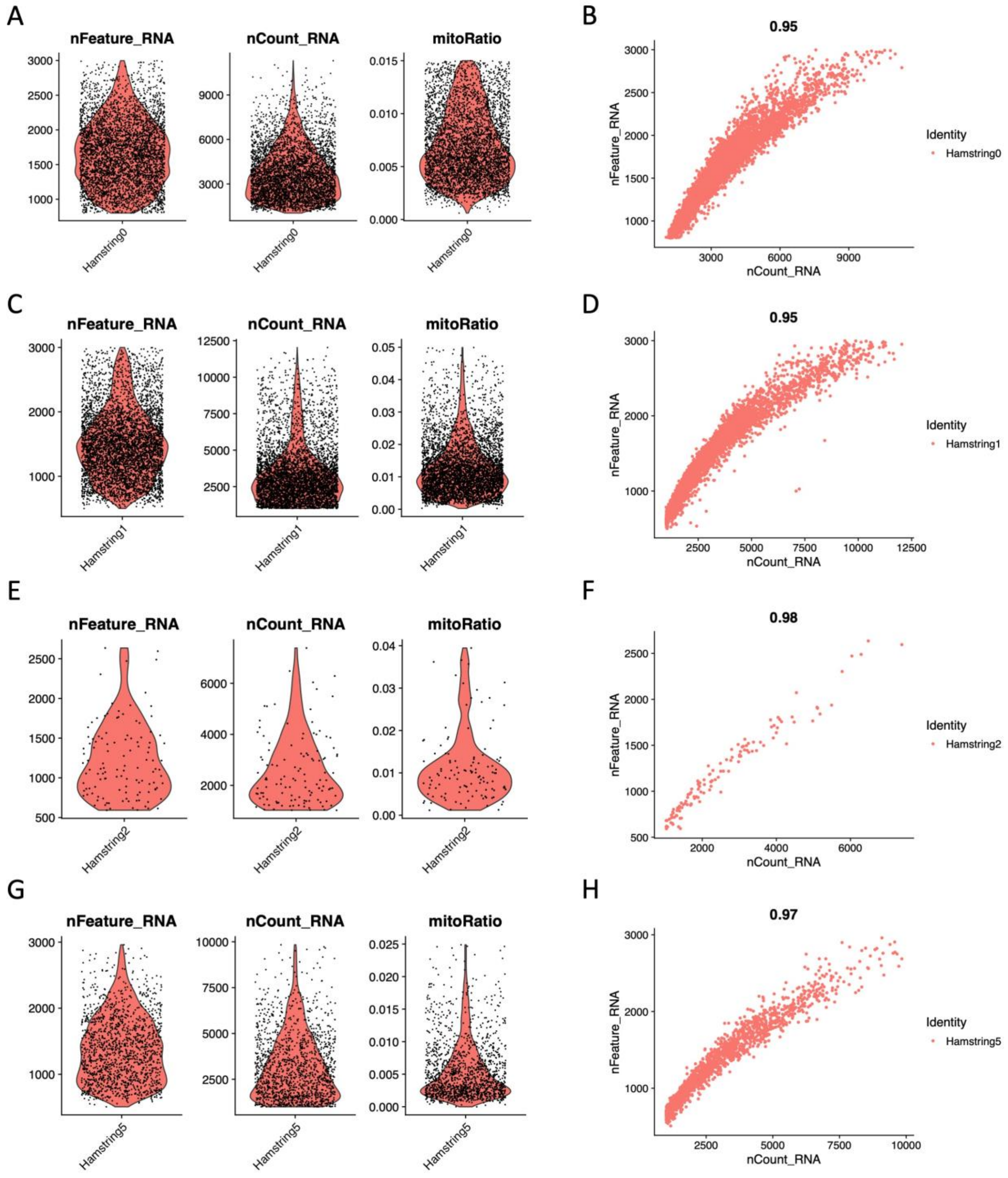


Supplementary Figure 1. Quality control of each dataset after filtering and removal of doublets and ambient RNA, including violinplots for the number of RNA features (nFeature_RNA), the number of RNA counts (nCount_RNA), and the mitochondrial ration (mitoRatio), as well as scatterplots of nFeature_RNA versus nCount_RNA. Datasets are hamstring0 (A-B), hamstring1 (C-D), hamstring2 (E-F), and hmastring5 (G-H).

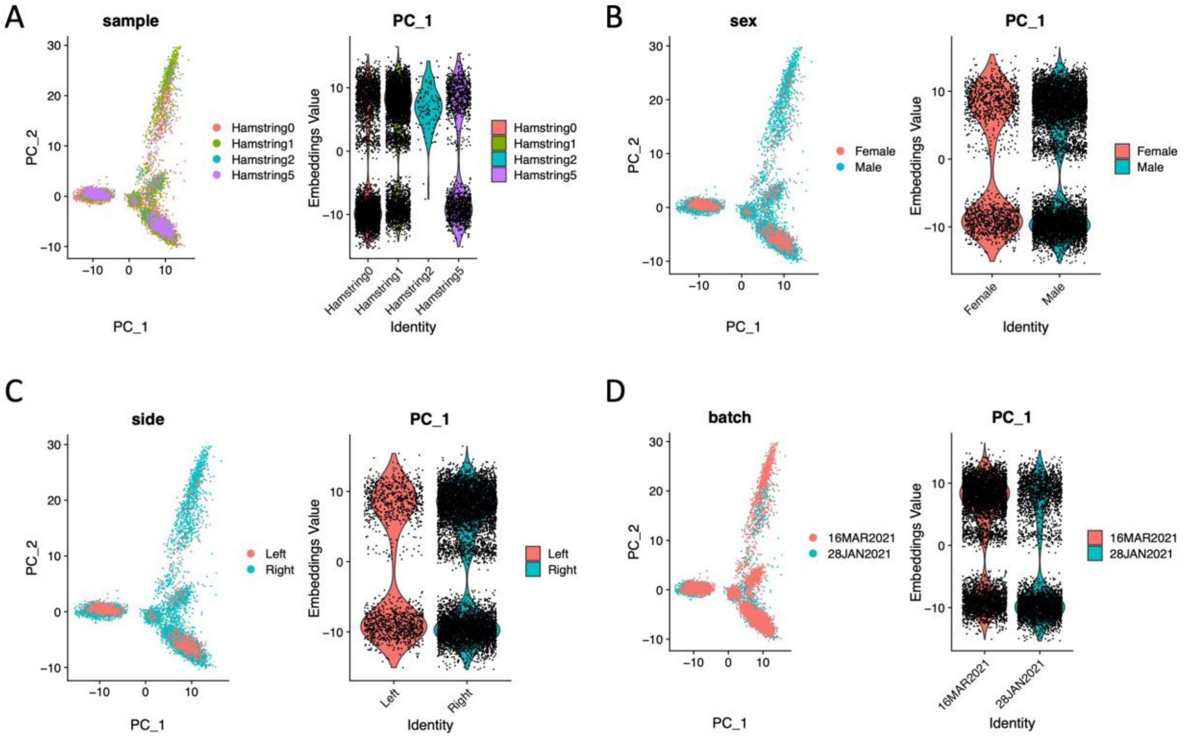


Supplementary Figure 2. The distribution of cells in the first two principle components (PC) based on the variables sample (A), sex (B), side of the body (C), and batch (D).

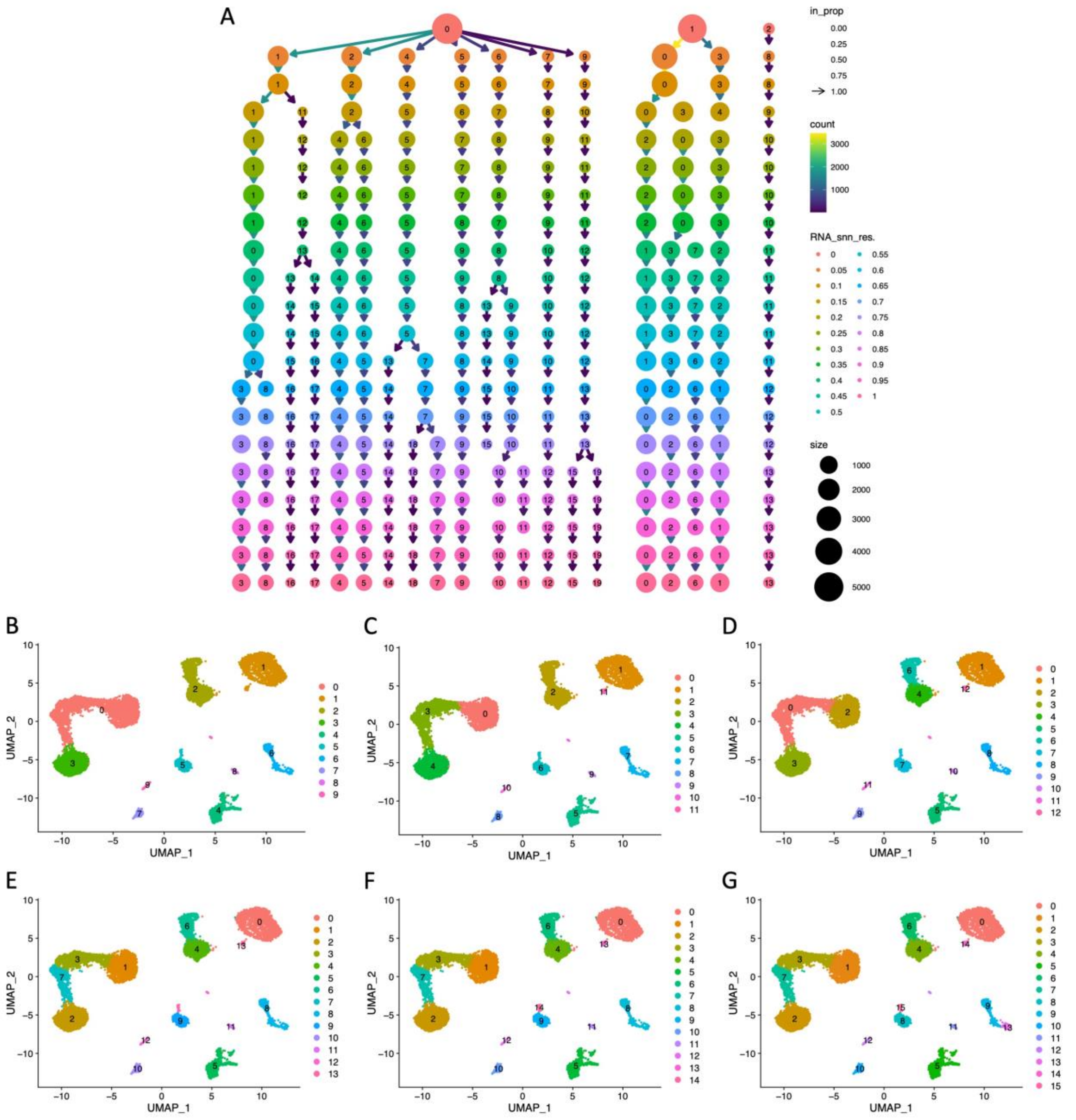

Supplementary Figure 3. Harmony clustering at 50 dimensions across different resolutions. (A) Clustree of resolutions 0 to 1 with 0.05 increments. (B-G) UMAPs from the first 6 resolutions in which changing in clustering occurs, namely resolution 0.05 (B), 0.15 (C), 0.20 (D), 0.40 (E), 0.45 (F), and 0.50 (G).

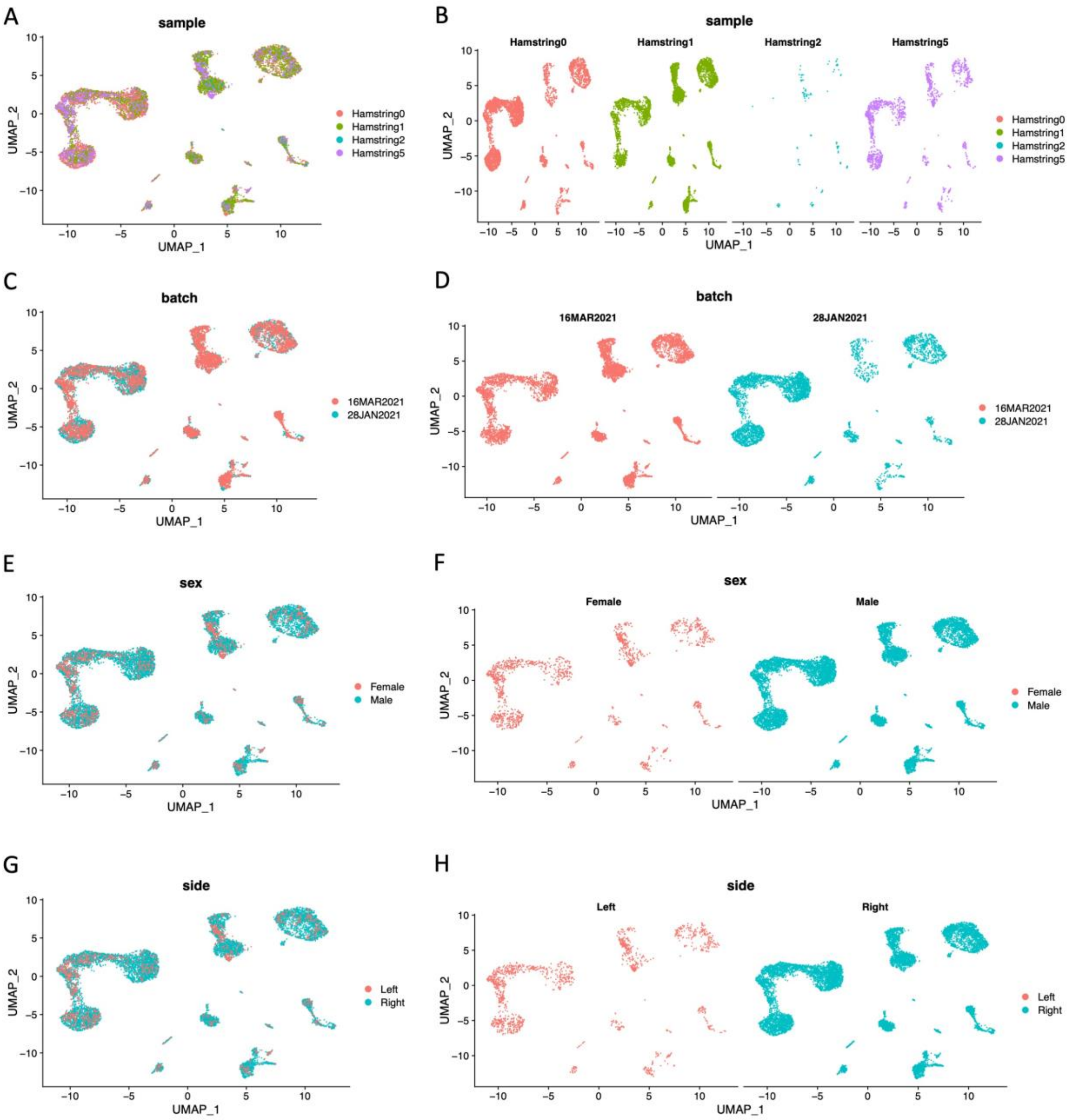


Supplementary Figure 4. The distribution of cells based on the variables sample (A-B), batch (C-D), sex (E-F), and side of the body (G-H).

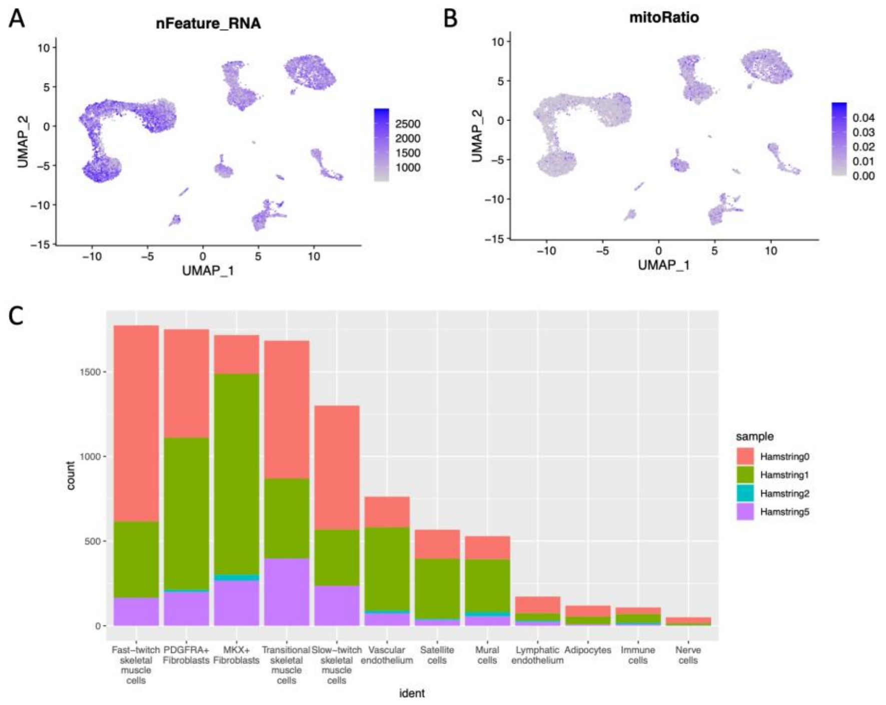


D

| Cluster name | Hamstring0 | Hamstring1 | Hamstring2 | Hamstring5 | Total |
|---|---|---|---|---|---|
| Fast-twitch skeletal muscle cells | 1,158 | 451 | 1 | 164 | 1,774 |
| PDGFRA+ Fibroblasts | 640 | 899 | 14 | 198 | 1,751 |
| MKX+ Fibroblasts | 229 | 1,187 | 36 | 265 | 1,717 |
| Transitional skeletal muscle cells | 815 | 472 | 1 | 396 | 1,684 |
| Slow-twitch skeletal muscle cells | 732 | 332 | 0 | 236 | 1,300 |
| Vascular endothelium | 181 | 492 | 16 | 73 | 762 |
| Satellite cells | 171 | 355 | 8 | 33 | 567 |
| Mural cells | 137 | 311 | 26 | 55 | 529 |
| Lymphatic endothelium | 98 | 44 | 8 | 22 | 172 |
| Adipocytes | 65 | 46 | 0 | 8 | 119 |
| Immune cells | 39 | 52 | 9 | 8 | 108 |
| Nerve cells | 36 | 12 | 1 | 1 | 50 |
| Total number of cells | 4,301 | 4,653 | 120 | 1,459 | 10,533 |

Supplementary Figure 5. Featureplots of the number of RNA features (A) and the mitochondrial ratio (B) of each cell plotted on the UMAP, as well as the number of cells each donor contributed to each identified cell type (C-D).

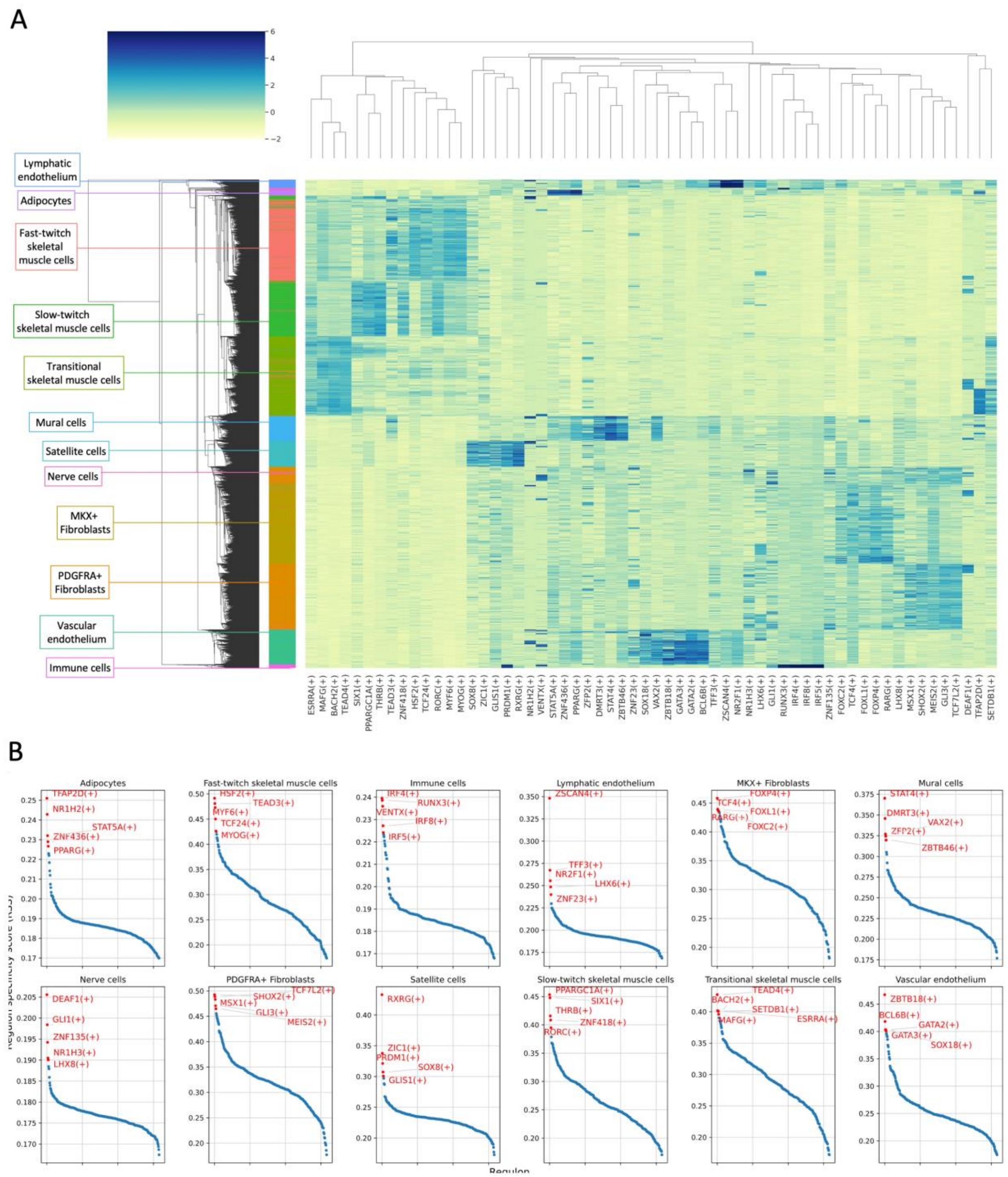


Supplementary Figure 6. SCENIC top 5 cluster-specific regulons. (A) Heatmap representing AUCell Z-scores for regulons with Top 5 RSS for each cluster. (B) Z-scores for each cluster, including the top 5 cluster-specific regulations highlighted in red.

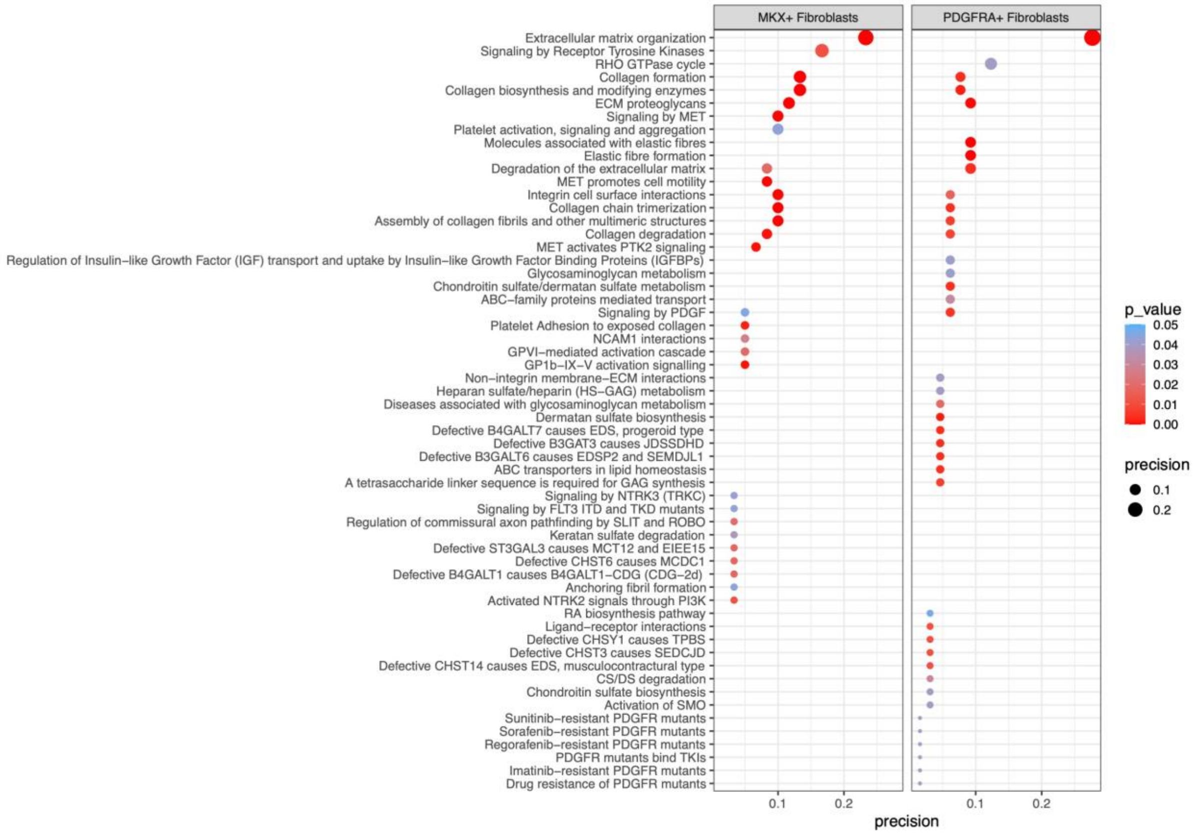


Supplementary Figure 7. Reactome pathways that are enriched in MKX+ fibroblasts and PDGFRA+ fibroblasts in healthy human hamstring tendon.

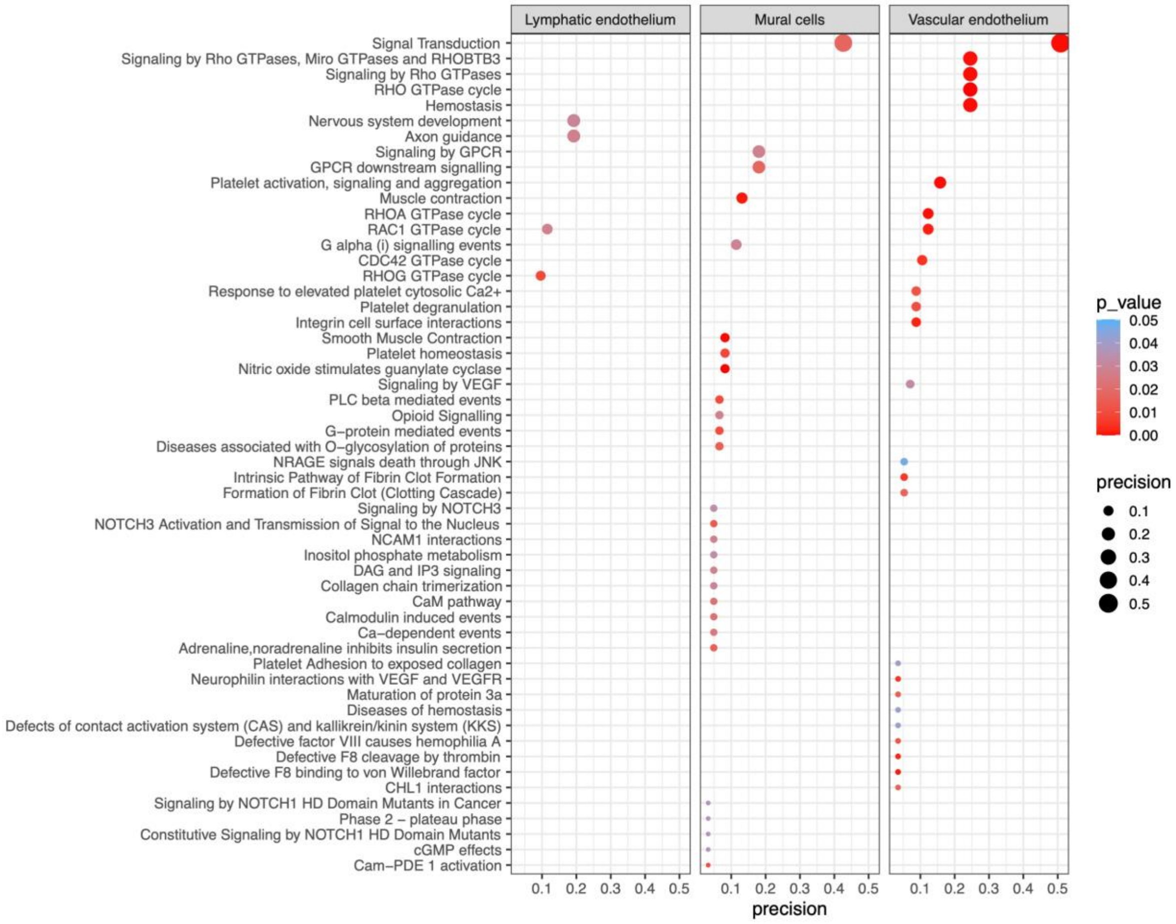


Supplementary Figure 8. Reactome pathways that are enriched in lymphatic endothelium, mural cells, and vascular endothelium in healthy human hamstring tendon.

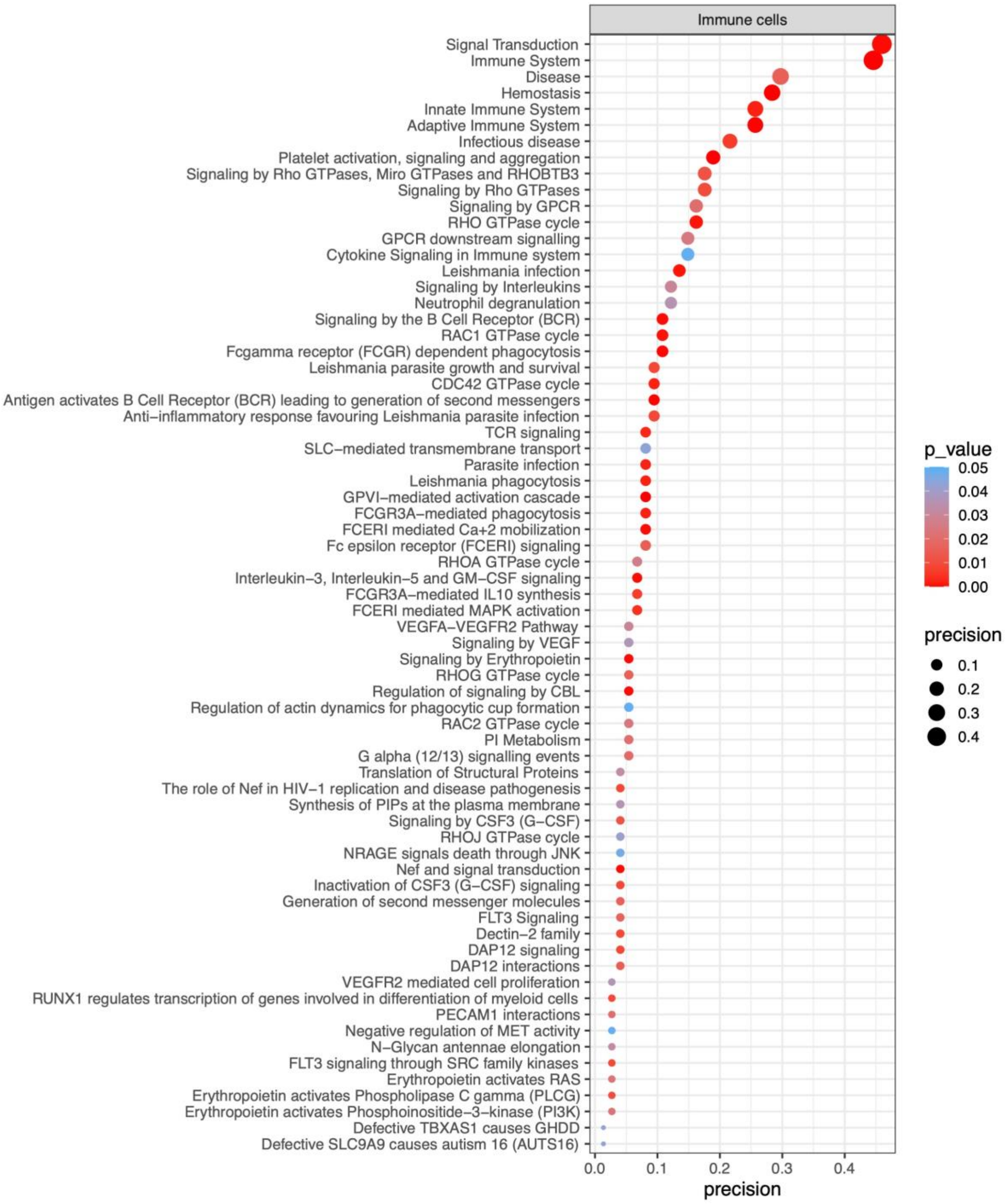


Supplementary Figure 9. Reactome pathways that are enriched in immune cells in healthy human hamstring tendon.

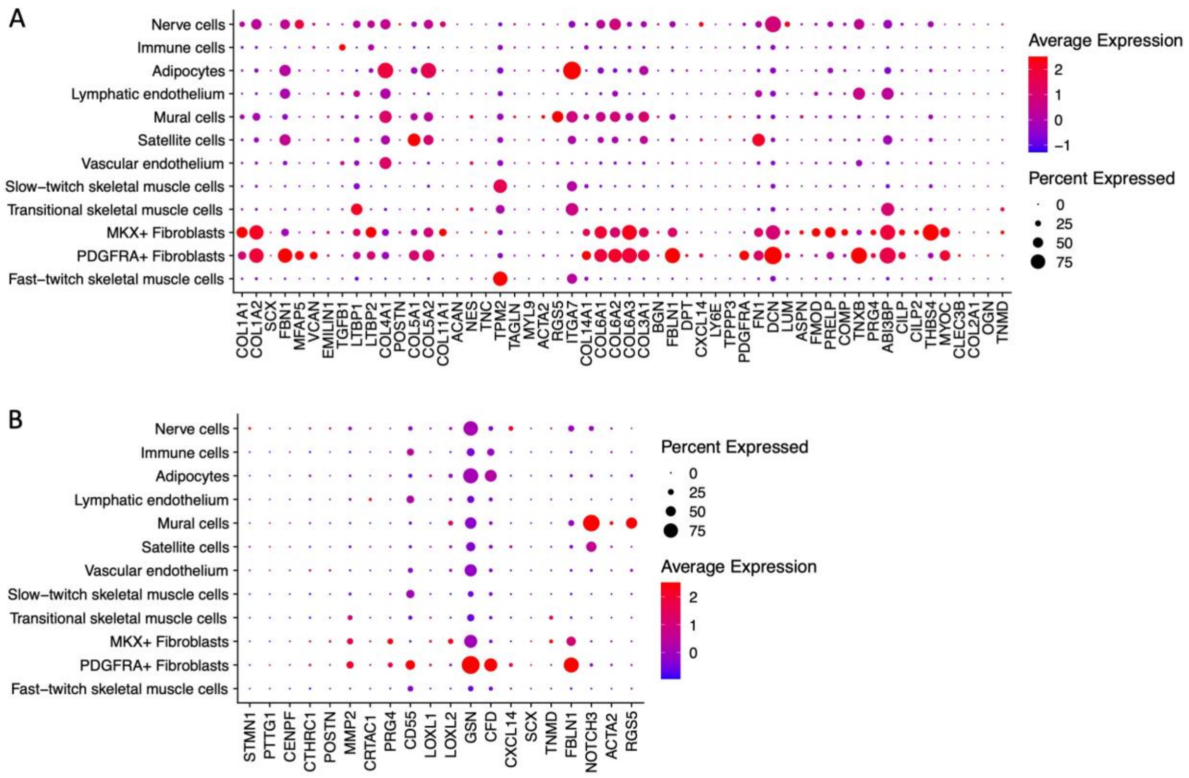


Supplementary Figure 10. Expression of markers used to identify specific tenocytes populations in the papers by (A) Kendal *et al.* (2020) and (B) Akbar *et al.* (2021).
